# Prediction of HLA genotypes from single-cell transcriptome data

**DOI:** 10.1101/2022.06.09.495569

**Authors:** Benjamin D Solomon, Hong Zheng, Laura W Dillon, Jason D. Goldman, Christopher S Hourigan, James R. Heath, Purvesh Khatri

## Abstract

The human leukocyte antigen (HLA) locus plays a central role in adaptive immune function and has significant clinical implications for tissue transplant compatibility and allelic disease associations. Studies using bulk-cell RNA sequencing have demonstrated that HLA transcription may be regulated in an allele-specific manner and single-cell RNA sequencing (scRNA-seq) has the potential to better characterize these expression patterns. However, quantification of allele-specific expression (ASE) for HLA loci requires sample-specific reference genotyping due to extensive polymorphism. While genotype prediction from bulk RNA sequencing is well described, the feasibility of predicting HLA genotypes directly from single-cell data is unknown. Here we evaluate and expand upon several computational HLA genotyping tools by comparing predictions from human single-cell data to gold-standard, molecular genotyping. The highest 2-field accuracy averaged across all loci was 76% by arcasHLA and increased to 86% using a composite model of multiple genotyping tools. We also developed a highly accurate model (AUC 0.93) for predicting *HLA-DRB345* copy number in order to improve genotyping accuracy of the *HLA-DRB* locus. Genotyping accuracy improved with read depth and was reproducible at repeat sampling. Using a metanalytic approach, we also show that HLA genotypes from PHLAT and OptiType can generate ASE ratios that are highly correlated (R^2^ = 0.8 and 0.94, respectively) with those derived from gold-standard genotyping.

**Highlights:**

- Benchmarking of HLA genotype prediction accuracy from 5’- and 3’-based single-cell RNA-seq data compared to molecular HLA genotyping
- Quantification of transcript coverage by 5’- and 3’-based single cell sequencing methods
- Accurate prediction of complex *HLA-DRB345* copy numbers using supervised learning
- Balancing accuracy and performance through composite HLA genotyping
- Meta-analytic approach to summarizing allele-specific expression of HLA genotypes at single-cell level

## INTRODUCTION

The human leukocyte antigen (HLA) locus is the most polymorphic region of the human genome. It encodes a wide range of important immune proteins, including the class I (HLA-A, -B, -C) and class II (HLA-DP, -DQ, - DR) major histocompatibility complex (MHC) proteins responsible for antigen presentation. The allelic diversity of the HLA locus contributes to the ability of MHC proteins to present a large range of possible peptides to lymphocytes and underlies the association of HLA genotypes with transplant compatibility and disease susceptibility [1].

HLA alleles are identified by four fields of increasing specificity, each with two digits. Together, the first two “low-resolution” fields define unique protein coding sequences, while the final two “high-resolution” fields denote synonymous -exonic and -intronic nucleic acid variation, respectively. The earliest HLA typing methods relied on low-resolution serologic assays that have largely been replaced by high-resolution molecular genotyping based on a variety of sequencing techniques. Clinical HLA genotyping typically utilizes site-specific amplicon sequencing of HLA loci, while research applications have increasingly focused on next generation sequencing (NGS) [2], [3].

HLA transcription is dynamically regulated throughout an inflammatory process and also represents a common immune escape mechanism. For example, reduced expression of MHC class 2 genes on monocytes correlates with mortality in septic shock [4] and multiple viral infections [5]. Regulation of HLA expression also extends to the level of individual alleles [6]. Temporal patterns in allele-specific expression (ASE) of certain *HLA-DQB1* alleles can discriminate their relative association with type 1 diabetes [7] and increased ASE imbalance of MHC class I genes was observed in colorectal [8] and other cancers [9]. Single-cell sequencing allows for fine resolution of ASE and such studies have demonstrated much of the transcriptome is mono-allelically expressed at a given point in time [10]–[12]. However, while methods for assessing single-cell ASE for HLA loci are available [13], they require high-resolution reference genotypes for accurate read mapping due to extensive HLA polymorphism, limiting their broader application to the increasingly large amount of publicly-available sequencing data.

Several computational tools exist to predict HLA genotypes from bulk RNA sequencing data and the accuracy of these methods have been robustly characterized [14]. However, it is unclear if these methods can be applied to data from single-cell RNA sequencing (scRNA-seq) experiments due to differences in sequencing chemistry and transcriptome coverage. One attempt to obtain genotypes from single cells observed that very few cells express HLA genes at a high enough level to generate genotype predictions for individual HLA loci [15].

Here, we show that data from commonly used scRNA-seq platforms can be condensed into subject-specific “pseudobulk” sequence files that can be used to predict HLA genotypes. Using five *in silico* HLA genotyping tools, we compare the accuracy of genotypes derived from scRNA-seq to gold standard molecular HLA genotyping obtained from the same individuals. We further expand on these methods to better predict complex *HLA-DRB345* genotypes and obtain maximal accuracy using a composite of the tested genotyping tools. We also show that even inaccurate genotypes from several tools can result in an accurate assessment of ASE in downstream use.

## METHODS

### HLA multiple sequence alignment and variation

HLA-allele sequences were obtained from IMGT/HLA version 3.42.0. All sequences without atypical expression suffixes (e.g. –N, -L, etc.) were included in multiple sequence alignment performed by DECIPHER/2.18.1 in R/4.0.4. Sequence variability was determined using 2-bit Shannon entropy based on nucleotide identity at each sequence position. Gap nucleotides were not factored into entropy calculations, as published HLA reference alleles often include only partial exon sequences

### Samples, sequencing, and gold standard HLA genotyping

Samples used for 5’ scRNA-seq have been described previously [16] as part of the ISB-Swedish INCOV study and include peripheral blood samples obtained from 157 patients at one or two time points. Samples were processed using 10X Genomics 5’ Chromium Single Cell Kits and sequenced using the Illumina Novaseq platform. Samples used for 3’ scRNA-seq and bulk RNA-seq have been previously described [17] as part of the NIH-HBM cohort and include bone marrow samples from 20 healthy patients. Samples were processed with using 10X Genomics Single Cell 3′ Solution Kits, TruSeq Stranded Total (bulk) RNA Sample Preparation Kits, or both. Libraries were sequenced on Illumina HiSeq 3000. For both sample cohorts, 3-field molecular HLA genotyping was performed by Scisco Genetics on peripheral blood aliquots or gDNA.

### FASTQ file preparation

For scRNA-seq samples, raw FASTQ files were demultiplexed into subject-specific FASTQ files utilizing UMI-tools/1.1.1. Quality control was performed with TrimGalore/0.6.5. Files were then mapped to the GRCh38 human reference genome with HISAT2/2.2.1 and indexed with SAMTOOLS/1.9. The ‘extract’ function of arcasHLA/0.2.0 was used to isolate chromosome 6 and unmapped reads to be used for downstream HLA mapping and genotyping.

### HLA genotype prediction

Genotyping by arcasHLA was performed with the ‘genotype’ function. The ‘parameters.p’ file in arcasHLA was modified to allow for genotyping of the HLA-DRB4 locus. Genotyping by PHLAT/1.1, OptiType/1.3.3, and HLAMiner/1.4 and scHLAcount/DEV were performed using default settings. When able to specify a user-defined HLA reference, IMGT/HLA version 3.42.0 was used.

### Prediction validity

For a given genotyper, success compares only those loci that a genotyper generated a prediction for to the gold standard and is represented by:

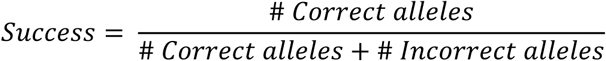

In contrast, accuracy reflects the validity of a given genotyper across all alleles in the gold standard, even those that the genotyper failed to generate a prediction for. This is analogous to Bray-Curtis similarity and is represented by:

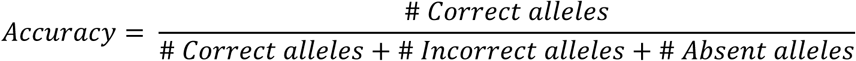

“Correct allele” indicates an allele prediction that matches the ground truth allele. Agreement between two predicted genotypes is a case of accuracy where “correct allele” indicates the two predicted genotypes are identical. Genotype accuracy and success were assessed independently at each HLA locus and thus only take values of 0.0, 0.5, or 1.0, reflecting 0, 1, or 2 correct alleles out of 2 possible alleles.

### Read depth analysis

For read subsampling, FASTQ files generated by arcasHLA extract were subsampled to 10%, 1%, and 0.1% of their original read count using seqtk/1.3. For cell subsampling, cellular barcodes were subsampled to 10%, 1%, and 0.5% of total sample barcodes. All reads corresponding to subsampled barcodes were isolated from FASTQ files generated by arcasHLA extract using BBMAP/38.90. Subsampled files were genotyped as described above. Linear modeling of genotyping accuracy/success by read/cell depth was performed with ordinary least squares regression on log-transformed read counts using base R/4.0.4.

### Statistical modeling

*HLA-DRB345* allele copy numbers were predicted from a K-nearest neighbor classifier using the ratio of *HLA-DRB3, -DRB4*, or -*DRB5* mapped reads to *HLA-DRB1* mapped reads as feature and the number of *HLA-DRB3, - DRB4, -DRB5* alleles identified by molecular genotyping as ground truth. Multi-class AUC was determined using the Hand & Till method [18]. Manhattan distance was used to quantify similarity between the allele count vectors {*-DRB3, -DRB4*, -*DRB5*} obtained from the kNN model and molecular genotyping. This distance was standardized to the maximum possible distance of 6, representing complete dissimilarity of the two alleles at each of the three loci. This value was then inverted to create an index where 1 represents complete similarity of all allele counts and 0 represents complete dissimilarity. Decision trees for composite HLA genotypes included presence/absence of a valid genotype from each genotyper, locus, and field level as input features and the identity of the highest accuracy genotyper for a given sample as ground truth. Both models were trained on 70% of the 5’ scRNA-seq samples with 10-fold cross validation and tested on the 30% hold-out set using Tidymodels/0.1.3 in R/4.0.4.

### Analysis of single-cell RNA-seq data

Gene expression analysis of 5’ scRNA-seq samples including pre-processing, integration, and cellular annotation was published previously [5]. Allele specific expression was determined by scHLAcount/DEV (commit 5ce7b2d) and incorporated into single cell data using Seurat.

### Allele-specific expression effect size

To prevent lower-confidence ASE ratios from cells with low read counts from biasing the overall sample ASE, we determined a summary effect size using the log-odds of each cell’s HLA expression ratio. For an individual cell, expected allele counts were calculated by summing the total number of reads between the two alleles, then distributing them equally between the two alleles. An odds ratio was obtained from the observed counts and this expected count, with the highest expressed allele as reference. To obtain a summary effect size across all cells, this odds ratio was converted to log-odds then weighted by the inverse variance of the log-odds. These weighted odds were summed in a random-effects model to find the effect size. Computation was performed using the R-package meta/4.19.0.

## RESULTS

### Single cell sequencing methods can produce sufficient sequence coverage to assess HLA sequence diversity

Compared to paired-end bulk RNA-seq methods, most high throughput scRNA-seq methods generate cDNA libraries that are enriched for either the 3’ or 5’ ends of transcripts. Given the extensive polymorphism of the HLA locus, incomplete sequence coverage could impair genotyping accuracy by excluding sequence variations outside of these enriched regions. This is particularly true for 3’-based protocols, as HLA diversity is concentrated in the 5’ region of each gene (**Figure 1A**).

**Figure 1:**
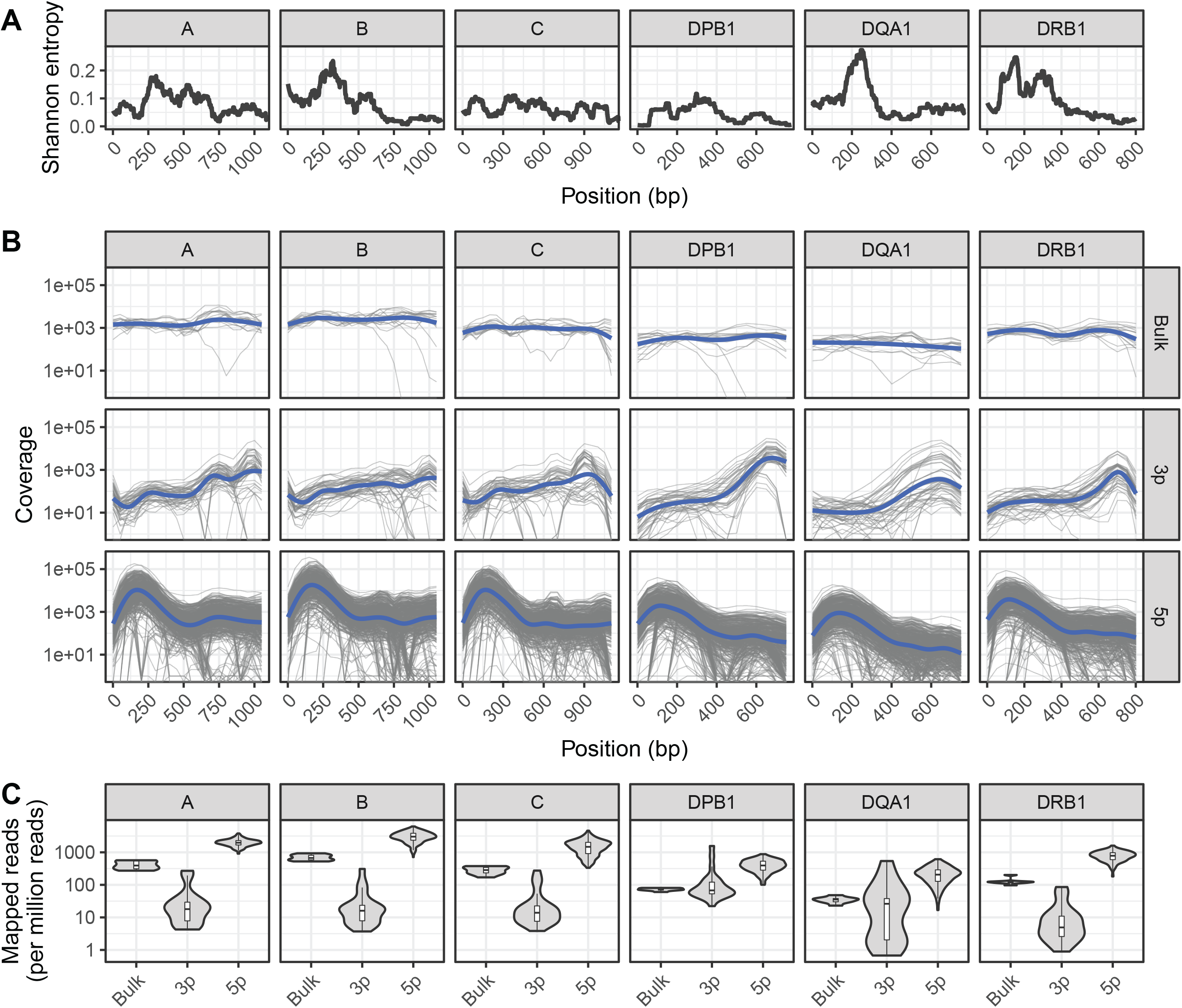
Sequencing coverage of HLA allelic diversity. A) Rolling (100bp) mean Shannon entropy for published allele sequences of indicated HLA loci. **B)** HLA coverage of reads mapped from bulk RNA-seq, 3’ (3p-based) scRNA-seq, and 5’ (5p)-based scRNA-seq. Grey lines represent individual samples, blue lines represent loess regression. **C)** HLA-mapped reads per million total reads from bulk RNA-seq, 3’ (3p)-based scRNA-seq, and 5’ (5p)-based scRNA-seq.

To quantify the relationship between HLA-sequence diversity and the positional bias of sequencing platforms, we compared the HLA loci coverage of 3’- and 5’-based scRNA-seq data, as well as bulk RNA-seq data. We selected two RNA-seq data sets with matched molecular HLA genotyping for this comparison and all downstream analyses. The first dataset, referred to as INCOV, included 5’-based scRNA-seq data from blood samples of 157 individuals taken at multiple time points [16]. The second dataset, referred to as NIH-HBM, included both 3’-based scRNA-seq and bulk RNA-seq data from healthy bone marrow donors [17].

Interestingly, while both 3’ and 5’ coverage bias was evident in the respective sequencing methods, both resulted in reads that could be mapped across the full extent of HLA loci (**Figure 1B**). In particular, coverage from 5’ based scRNA-seq data was comparable or better than that of bulk RNA-seq across the full extent of most HLA loci. This was further reflected in the overall mapping efficiency of reads in each HLA loci, as the median number of reads aligned to HLA loci from 5’-based scRNA-seq methods were an order of magnitude greater than both bulk RNA-seq and 3’-base RNA-seq data (**Figure 1C**).

### scRNA-seq pseudobulk data results in few HLA genotyping failures

Darby *et al*. previously demonstrated that individual cells do not express HLA transcripts at a sufficient level for reliable and complete genotyping at the single cell level [15]. Therefore, we sought to achieve sufficient read coverage at HLA loci by pooling reads from all cells of an individual sample into a “pseudobulk” data set. As most single-cell methods generate single-end reads, we identified 4 compatible HLA genotyping tools: arcasHLA [19], HLAminer [20], PHLAT [21], and OptiType [22]. Though primarily used to quantify ASE at HLA loci, we also compared predictions from the genotyping function of scHLAcount [13]. To control and minimize the variability resulting from the different global alignment strategies incorporated into each of these tools, we first performed common global alignment with HISAT2 then isolated chromosome 6 reads (containing the HLA locus) and unmapped reads to use as a starting point for each genotyper.

Overall, most tools generated two allele predictions for the first two HLA fields, while predictions at field 3 had a higher rate of failure (**Figure 2A**). Notably, arcasHLA and PHLAT made complete 2-allele predictions across all loci, with the exception of *HLA-DPA1* and *HLA-DPB1* which PHLAT does not assess. scHLAcount was a notable exception, typically producing only a single allele prediction.

**Figure 2:**
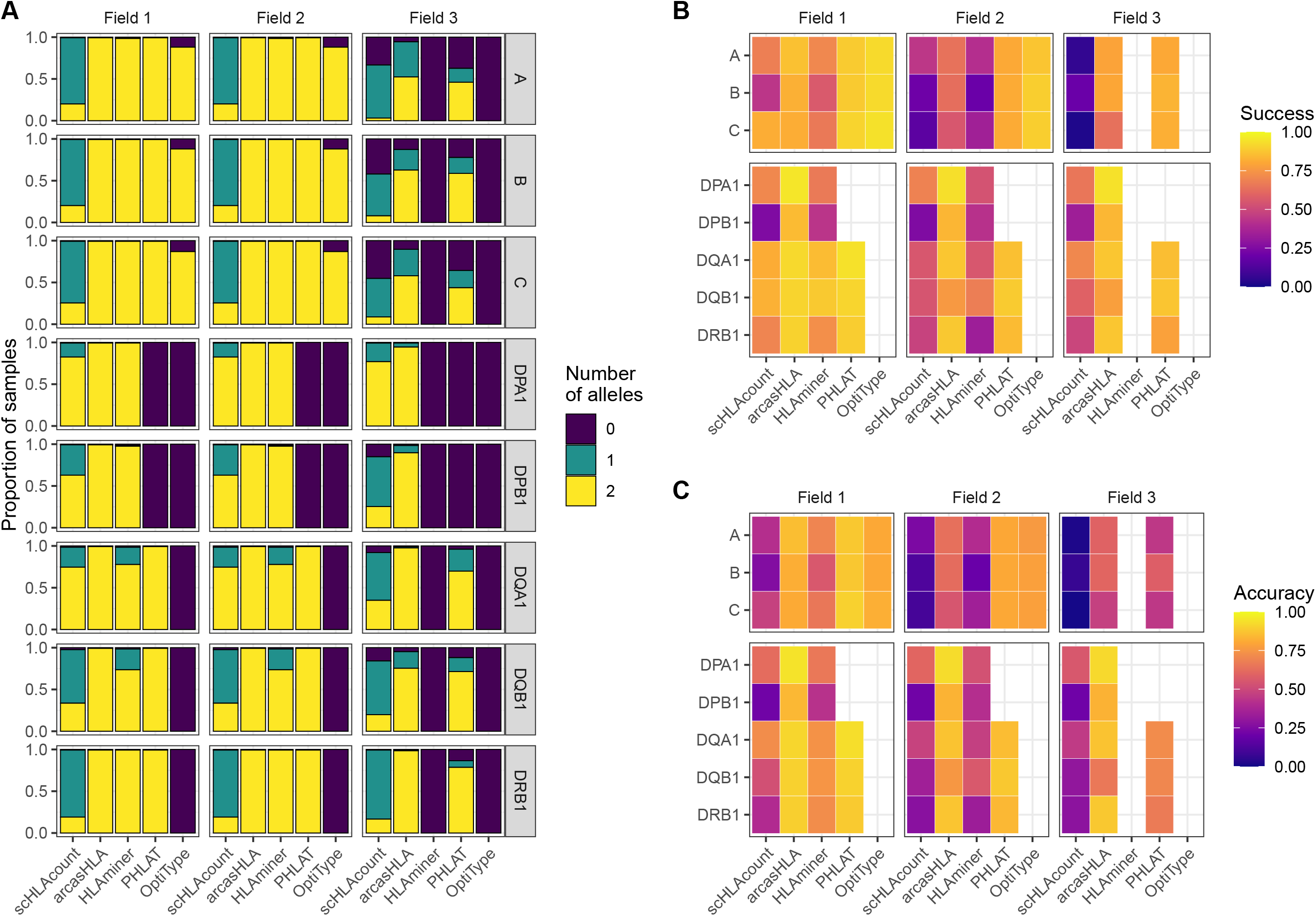
Accuracy of HLA genotype predictions. **A)** Relative proportion of samples with 0, 1, or 2 allele predictions. **B)** Mean success and **C)** mean accuracy of predicted genotypes compared to molecular genotyping. Empty heatmap tiles represent genotype paramaters not assessed by a given genotyper.

### Genotyping accuracy varies by loci, genotyper, and sequencing direction

We compared predicted HLA genotypes to ground-truth sample-matched molecular genotyping using two metrics previously utilized by Bauer *et al* [14]. “Success” describes the proportional match of a predicted genotype with the ground truth genotype but ignores missing predictions. Conversely, “accuracy” assesses how likely a genotyper is to predict the complete, correct genotype by penalizing failed genotype predictions.

We focused primarily on 5’-based scRNA-seq INCOV data due to its greater transcript coverage. For class 1 genes, OptiType had the highest success at fields 1 and 2, though lower accuracy compared to PHLAT due to occasional prediction failures of OptiType (**Figure 2B-C, Table S1 & S2**). aracasHLA had moderate success and accuracy for class 1 genes (55% - 64% at field 2), but had substantially higher accuracy for class 2 genes (75% - 93% at field 2). Its accuracy for class 2 genes and was only surpassed by PHLAT at *HLA-DQB1*. Notably, HLAMiner had the lowest success and accuracy compared to the other genotyping tools, with an average field 2 accuracy of less than 50% across all HLA loci. The genotyping function of scHLAcount also had poor accuracy. We excluded scHLAcount from further analysis due to its low accuracy and high rate of incomplete genotype predictions. This variation in accuracy did not result from bias of particular genotypes for specific alleles or samples (**Figure S2A-B**).

3’-based scRNA-seq NIH-HBM data resulted in lower accuracy than 5’-based data. The average 2-field MHC class 2 accuracy for arcasHLA ranged from 32% - 95% (**Figure S1A, Table S3**). Interestingly, MHC class 1 accuracy appeared less affected by the reduced sequence coverage compared to MHC class 2 accuracy. By comparison, bulk RNA-sequencing frequently resulted in excellent accuracy (**Figure S1B, Table S4**) such as average 2-field ranging from 88%-100% across all HLA loci for arcasHLA.

### Increasing accuracy of HLA-DRB345 predictions

The *HLA-DRB1* gene can occur alone or in close linkage disequilibrium with one of three functional paralogs, *HLA-DRB3*, -*DRB4*, or -*DRB5*, and pairing of *HLA-DRB1* and *HLA-DRB345* on each homologous chromosome occurs independently [23]. While arcasHLA and HLAminer can predict *HLA-DRB345* identity, neither accounts for copy number, frequently resulting in biologically invalid *HLA-DRB345* predictions.

To address this, we sought to predict the number of *HLA-DRB3 -DRB4*, and -*DRB5* copies prior to assessing allele accuracy. Zhang *et al*. previously showed that copy numbers could be predicted from targeted sequencing using the ratio of *HLA-DRB345* reads to *HLA-DRB1* reads [24]. Using a similar approach, we trained a K-nearest neighbor (kNN) classifier to predict the number of *HLA-DBR3, -DRB4*, and –*DRB5* alleles based on their relative read abundance (**Figure S3A**). We chose kNN for its low complexity and minimal assumptions when applied to multiclass modeling. We used 70% of the 5’-based scRNA-seq INCOV data to train the kNN-based classifier. The model was highly accurate when applied to a hold out subset (AUROC = 0.97; **Figure 3A**). This model generalized well to 3’-based scRNA-seq and bulk RNA-seq data from NIH-HBM, with an AUC of 0.79 and 1.0, respectively (**Figure S3D & S3F**).

**Figure 3:**
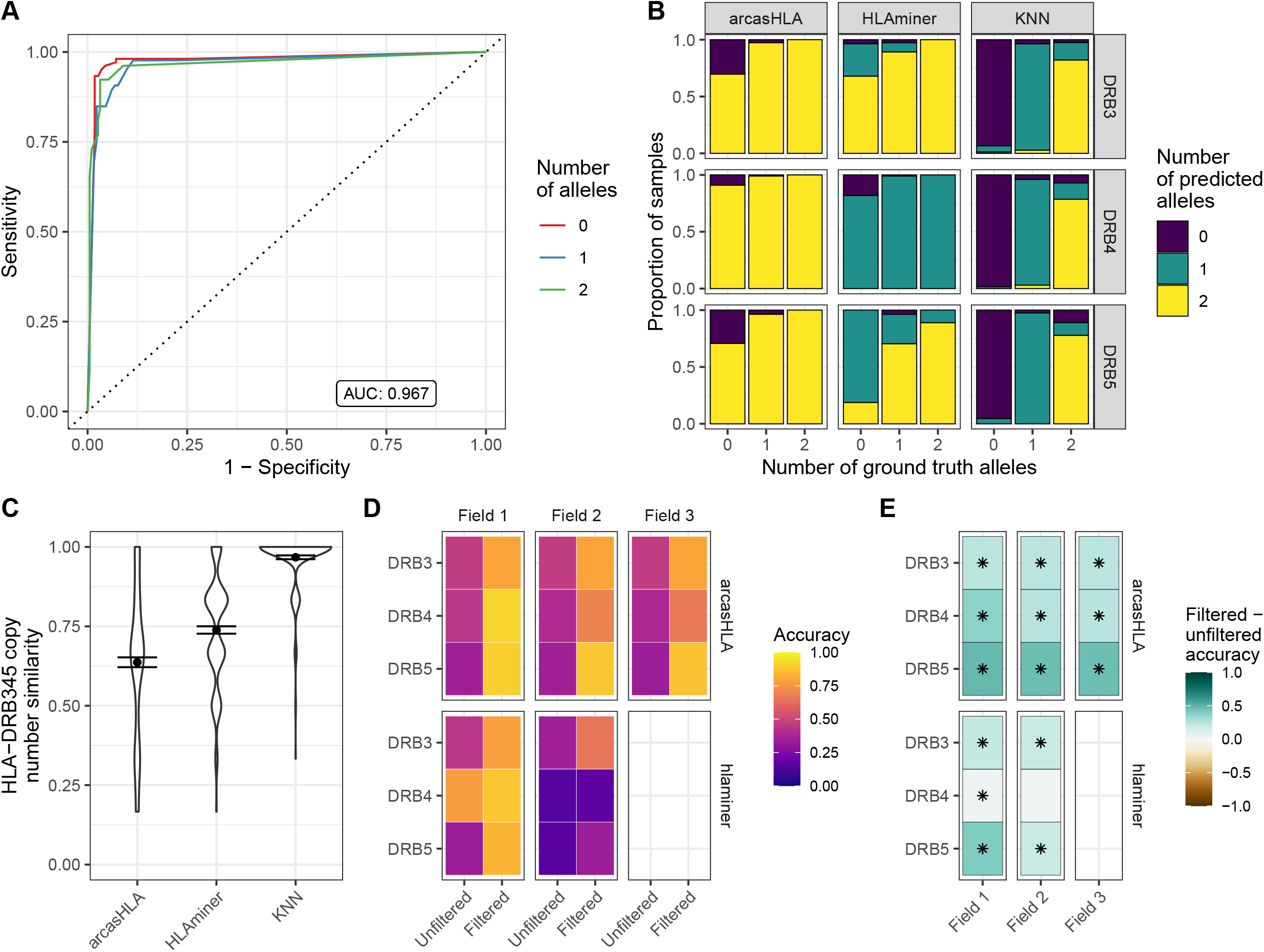
Prediction of HLA-DRB345 genotypes. **A)** ROC for HLA-DRB345 copy number classifier by kNN. Tuned on 70% test set with 10-fold cross validation. AUC represents performance on 30% hold-out test set. **B)** Relative proportion of samples with ground truth HLA-DRB3, -DRB4, and -DRB5 copy numbers of 0, 1, or 2 with corresponding predicted copy numbers by arcasHLA, HLAminer, or the kNN classifier applied to all samples. **C)** Similarity between the set of ground truth and predicted HLA-DRB3, -DRB4, and -DRB5 copy numbers for each sample. Similarity represents 1 - maximum-standardized Manhattan distance. Error bars represent standard error of the mean. **D)** Mean accuracy of HLA-DRB345 genotype predictions when unfiltered or filtered by kNN classifier. **E)** Difference in filtered and unfiltered mean accuracy. Asterisks represents an FDR-adjusted p-value ≤ 0.05 by Wilcoxon rank sum test.

Prior to application of the kNN classifier, copy numbers associated with genotype predictions were notably discordant from those reflected in ground truth genotypes. In samples with no HLA-DRB3 copies per molecular genotyping, arcasHLA and HLAminer correctly predicted an HLA-DRB3 copy number of zero in only 30% and 3% of these samples, respectively (**Figure 3B**). By comparison, the kNN classifier correctly identified 93% of these zero HLA-DRB3 copy number samples. When similarity to ground truth copy numbers was summarized across all HLA-DRB345 loci, the kNN classifier had a mean standardized similarity of 0.97, compared to 0.64 and 0.74 for arcasHLA and HLAminer, respectively (**Figure 3C**).

Using these copy number predictions, we filtered the *HLA-DRB345* genotypes from arcasHLA and HLAminer to their top *n* alleles, where *n* is the number of copies predicted by our kNN classifier. This significantly improved *HLA-DRB345* accuracy, indicative of a high frequency of false positive predictions in unfiltered genotypes. Accuracy ranges for 2-field arcasHLA predictions increased from 35%-46% using unfiltered samples to 66%-88% with filtered samples. (**Figure 3D-E, Table S5**). As with other loci, arcasHLA significantly outperformed HLAminer.

### Genotype predictions from scRNA data are precise

While HLA genotypes are static, differences in HLA gene expression levels might affect genotype predictions when based on scRNA-seq data. The 5’-based scRNA-seq dataset contained samples from individuals obtained at two time points, allowing us to assess prediction reproducibility. Both time points had similar prediction patterns, with no significant variation in accuracy when compared to molecular HLA genotyping (**Figure 4A, Table S6)**. In the case of arcasHLA, the difference in average 2-field accuracy between the two time points ranged from 0%-12%.

**Figure 4:**
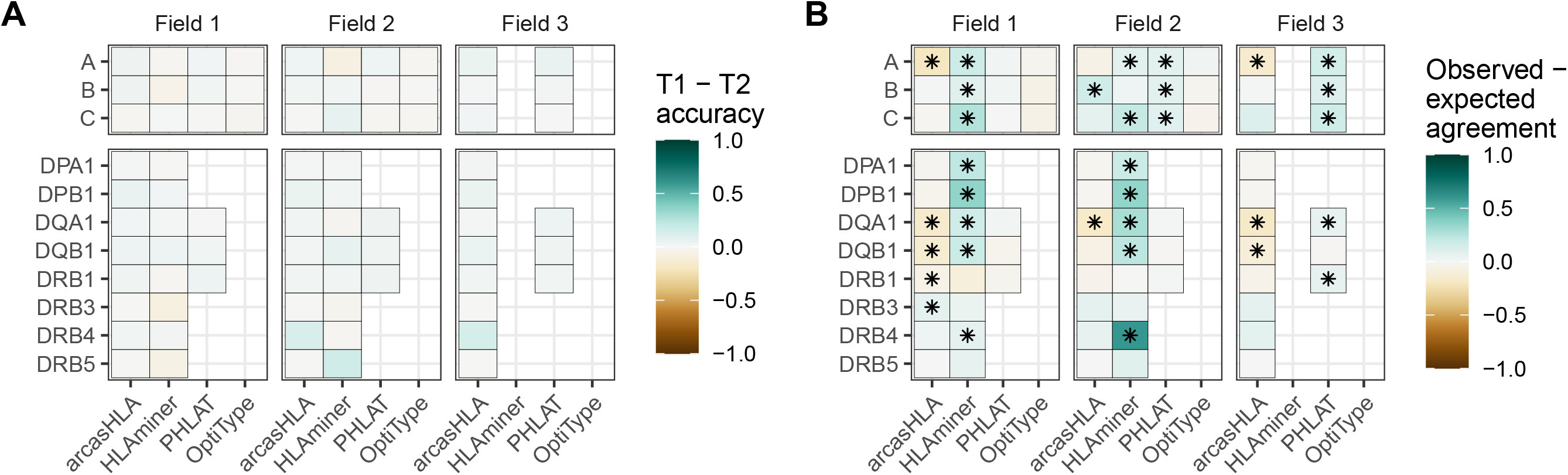
Reproducibility of predicted HLA genotypes. **A)** Difference in mean accuracy between genotype predictions at time point 1 (T1) and time point 2 (T2). **B)** Mean difference between the expected agreement of T1 and T2 genotype predictions and their observed agreement. Asterisks represents an FDR-adjusted p-value ≤ 0.05 by Wilcoxon rank sum test.

Next, we directly compared how well the genotypes from both time points agreed with one another. An expected agreement between time points was derived from the product of their respective accuracy values. For arcasHLA and OptiType there was no significant difference between expected and observed agreement at the 2-field level with the exception of arcasHLA at *HLA-DQA1*, which showed lower observed agreement (**Figure 4B**). Alternatively, observed 2-field agreement was either equivalent to or significantly higher than expected agreement for HLAminer and PHLAT across all HLA loci.

### Increased read depth improves genotyping accuracy

Previous studies have shown conflicting results regarding the effect of read depth on the accuracy of HLA genotyping from NGS data [14], [25]. We evaluated this relationship by assessing genotype accuracy from multiple levels of subsampled reads. Overall, accuracy increased with read depth for most genotypers and loci (**Figure 5**). OptiType accuracy was relatively constant over read depth due in part to genotyping failures at higher read counts (**Figure S4**). Conversely, the positive trend between read depth and accuracy was most prominent with arcasHLA, accentuated by its minimum read parameter that excludes genotype predictions for loci with a total number of mapped reads below a specified threshold.

**Figure 5:**
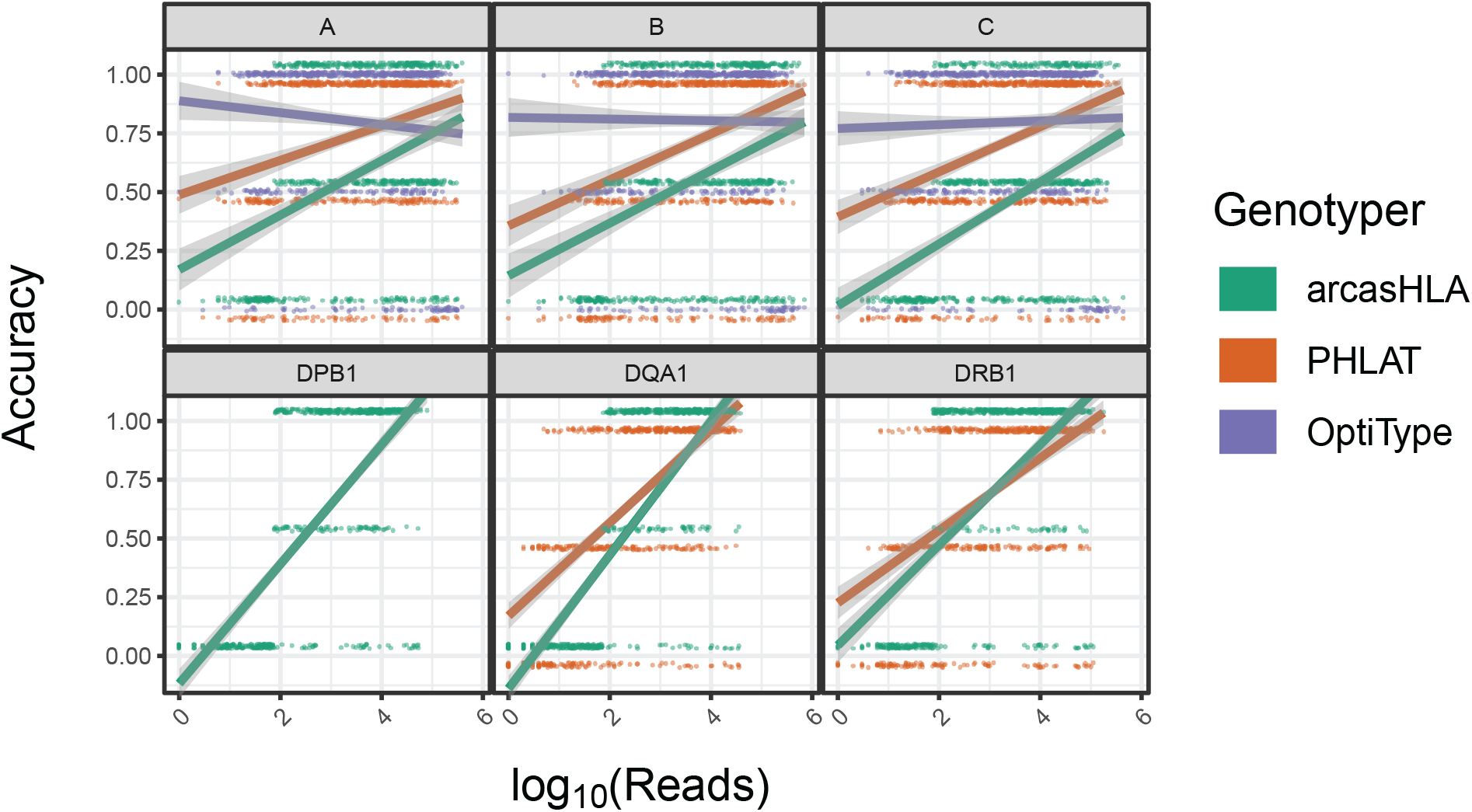
Effect of sequence sampling depth on HLA predictions. 2-field accuracy based on number of reads aligned to the indicated locus. Raw sequencing data sampled to 100%, 10%, 1%, and 0.1% of original sample read total. Points represent individual samples offset from {0, 0.5, 1} to distinguish genotypers. Trend lines represent linear regression using log transformed reads with ribbon representing 95% CI.

### Composite genotypes increase overall accuracy

Since no single genotyper produced the most accurate predictions across all samples and HLA loci, we sought to determine if combing multiple predictions could increase accuracy. The genotypers tested in our study have significant variation in run time ranging from a median runtime of 1.6 minutes per sample for arcasHLA to a median runtime of 183 minutes per sample for HLAMiner (**Figure 6A**). As such, when combining genotyper predictions, we also sought to balance any gain in accuracy with the increased processing time needed to run multiple tools.

**Figure 6:**
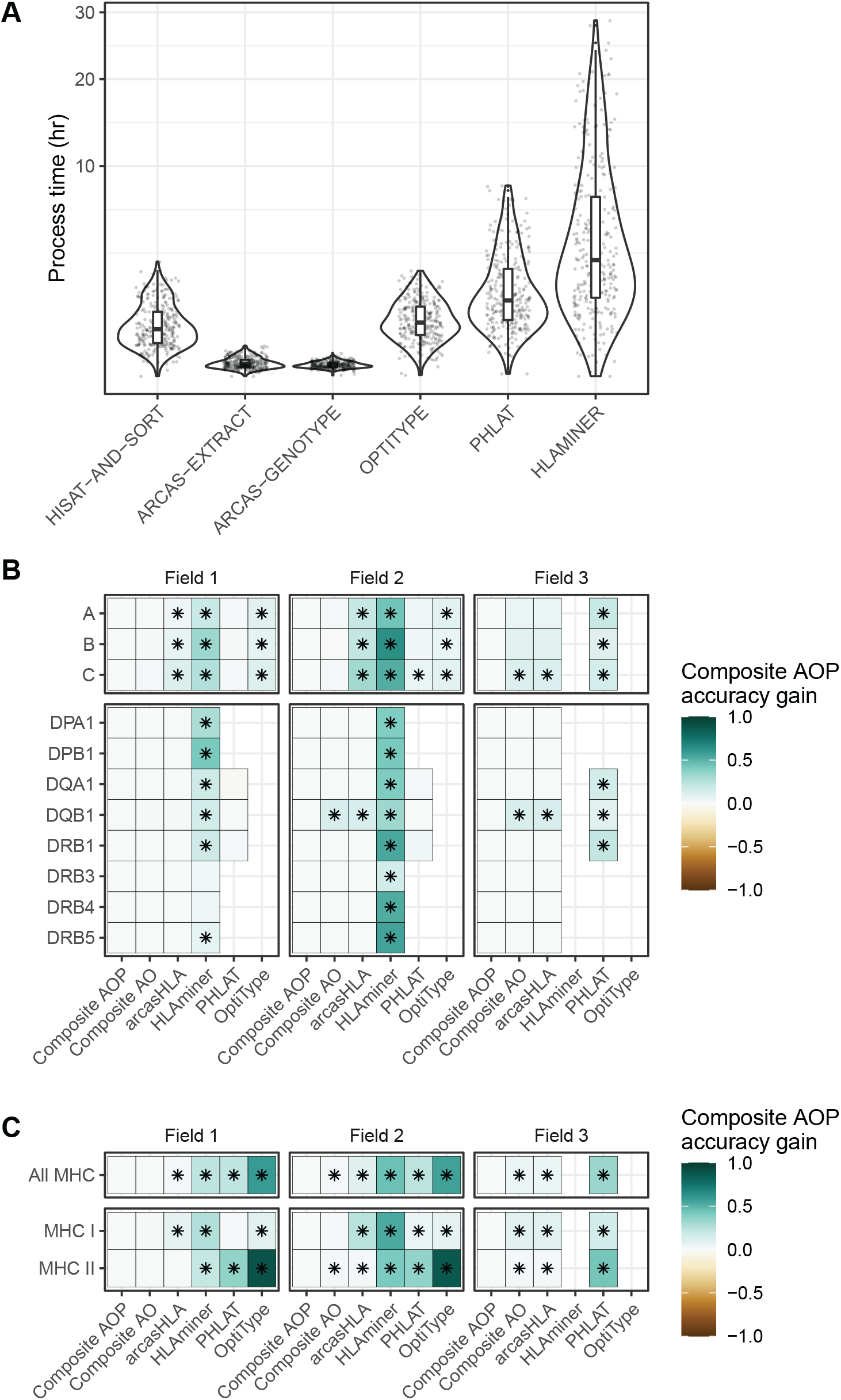
Composite HLA prediction accuracy. **A)** Runtime of major steps in genotyping pipeline. Y-axis distributed along square root scale. **B-C)** Difference in mean accuracy between AOP composite and indicated alternative genotyping methods across **B)** individual loci and **C)** loci classes. (AOP) arcasHLA, OptiType, PHLAT, (AO) arcasHLA, OptiType. Asterisks represents an FDR-adjusted p-value ≤ 0.05 by Wilcoxon rank sum test.

We trained a decision tree classifier to determine the most accurate genotyper based on (1) the set of all genotypers with successful predictions for an individual sample, (2) the HLA locus, and (3) the field level. We trained two trees, one that incorporated arcasHLA, OptiType, and PHALT (AOP) and one with only arcasHLA and OptiType (AO) (**Figure S6A-B**). HLAminer was excluded entirely due to its long run time and overall low accuracy. When tested on a 30% hold out dataset, both models were highly accurate at identifying the optimal genotyper for a given sample (AOP AUC 0.84, AO AUC 0.93).

We then created a composite HLA genotype for each individual based on the genotype prediction from the optimal genotyper at each locus and allele field level. Composite accuracy was consistently higher than accuracy from individual genotypers (**Figure 6C-B, Table S7**). 2-field accuracy from the AOP composite ranged from 84%-86% for MHC class 1 loci and 69%-93% for MHC class 2 loci. Notably, the accuracy of predictions from the AO composite were not significantly different from those obtained by the AOP composite with the exception of *HLA-DQB1*, suggesting that a near optimal prediction can be obtained from the combination of only arcasHLA and OptiType.

### Inaccurate genotype predictions can result in accurate determination of HLA allele-specific expression

Single-cell studies provide enhanced resolution of allele-specific transcriptional regulation, though analysis of ASE at HLA loci is complicated by the effect of HLA polymorphism on read mapping. This can be overcome by supplying sample-specific HLA references to tools like scHLAcount [13], though it is unclear how much allele-mistyping affects downstream determination of ASE ratios. To address this question, we evaluated how variation in accuracy from different HLA genotypers affected ASE.

In a representative sample **(Figure 7A**), inaccurate genotypes resulted in broadly different patterns of ASE (**Figure 7B**). To quantify this apparent difference in ASE across cells, we used a meta-analytic approach to summarize ASE ratios. When focusing on CD14+ monocytes from a representative sample, this approach demonstrated that the genotyping inaccuracy associated with HLAminer resulted in a notable deviation in the ASE log-odds ratio from that obtained using the ground truth genotype (**Figure 7C**).

**Figure 7:**
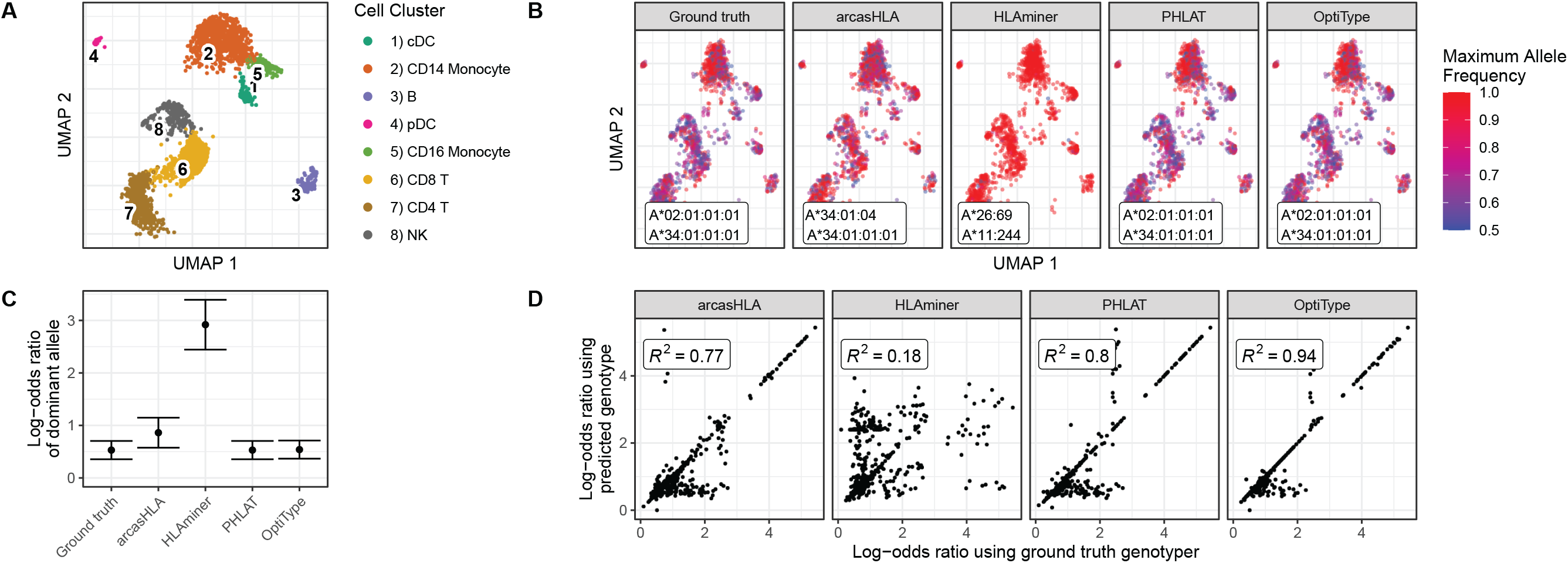
Effect of prediction accuracy on analysis of allele-specific HLA-A expression. **(A-C)** Representative sample from 5’ data set. **A)** UMAP projection of single-cell transcriptome data, annotated as previously described [4] **B)** UMAP plots colored by cell-specific expression frequency of the most highly expressed HLA allele determined by scHLAcount. The reference genotype from each genotyper used by scHLAcount is annotated. **C)** Summary log-odds ratios of dominant HLA-A allele from all cells in cDC cluster determined by random effects model. Error bars represent summary standard error. **D)** Correlation of ground truth- and genotyper-derived HLA-A allele log-odds ratio from all samples and cell types.

When comparing ground truth- and genotyper-derived ASE log-odds ratios across all samples and cell types, PHLAT and OptiType resulted in the greatest correlation with ground truth-derived ASE, with R^2^ values of 0.8 and 0.94, respectively (**Figure 7D**). Interestingly, this high level of correlation occurred even in samples where both allele predictions were incorrect, though not when only a single allele prediction was correct (**Figure S7**). This suggests that, even when inaccurate, these tools predict genotypes with sufficient sequence similarity to the ground truth genotype to allow for similar proportional mapping of reads to the two reference alleles.

## DISCUSSION

In this study, we describe the feasibility and best practices for obtaining HLA genotypes from scRNA-seq data. Using bulk RNA-seq data as a baseline, we found that our evaluation pipeline generated similar accuracy results to those seen in other benchmarking studies [14], [19]. By comparison, sequencing from 5’-based single-end scRNA-seq data was moderately accurate with average accuracy across all loci for arcasHLA of 76% and 74% for 2-field and 3-field genotypes, respectively. This accuracy could be increased by composite predictions assembled using a decision tree model from multiple genotypers, resulting in an average accuracy of 86% and 78% for 2-field and 3-field genotypes, respectively.

In order to obtain accurate genotype predictions of highly polymorphic regions like the HLA locus, full transcript coverage is critical to ensure all underlying sequence diversity is incorporated into predictions. We observed that 5’-based scRNA-seq methods provides excellent read coverage across the entire length of HLA transcripts compared to 3’-based scRNA-seq coverage, which is more restricted and enriched for regions with lower HLA polymorphism. Both protocols utilize a template switch reaction, though the proportion of the poly-A tail binding region of reverse-transcription primer used in the 5’-based protocol is nearly double that of the primer in the 3’-based protocol, possibly contributing to the difference in coverage we observe. The observations that transcript coverage from 5’-based scRNA-seq is sufficient to genotype highly polymorphic HLA loci, suggests that this platform could also reliably quantify other areas of small sequence variation such as single nucleotide polymorphisms.

Our observation that genotyping accuracy improves with increased locus coverage contrasts with that of Bauer *et al* [14]. Several differences may account for this. First, the bulk RNA-seq data analyzed previously is likely more robust to positional HLA sequence variation than the single-end scRNA-seq data assessed here. As such, more read depth may be necessary to overcome the relative loss of allele-identifying sequence information in single-ended libraries compared to paired-end libraries. In addition, our subsampling approach ensured that our analysis would test genotypers at the lowest limits of starting material. Without subsampling, the more limited distribution of coverage may fall well within the optimal range of sensitivity for the genotyping tools tested.

At 86% 2-field accuracy, even optimized composite predictions for 5’ scRNA-seq data are unlikely to be useful for clinical genotyping. However, the primary novel application of HLA typing to single-cell data is its research use in assessing ASE. For instance, using bulk RNA-seq, Liu *et al*. demonstrated that colorectal cancer samples were associated with greater skewing of HLA ASE ratios. Similar to studies that demonstrated an association between loss-of-HLA-heterozygosity with multiple tumor types and failure of therapeutic checkpoint blockade [26], this skewing away from equal biallelic HLA expression could represent an immunological escape mechanism that functions to reduce the diversity of possible tumor peptide-antigens displayed by MHC molecules. Interestingly, in our study, we show that even inaccurate genotype predictions from arcasHLA, PHLAT, and OptiType can result in ASE ratios that correlate highly with those derived from ground truth genotypes. This suggests that these inaccurate predictions still identify alleles with sufficient sequence similarity to the true alleles that proportional read mapping between the homologous chromosomes is maintained, resulting in relatively accurate assessments of ASE.

However, molecular HLA genotyping still represents the gold standard for accuracy and is readily available through commercial laboratories and kits. Yet, it is not always feasible to perform these assays for single-cell experiments. As scRNA-seq methods are limited by cost and cell number, building robust data sets often requires integration of published data, for which prior HLA genotyping or remaining biological samples may not be available. Moreover, particularly in the case of clinical studies, limited sample material may preclude the use of multiple analytical platforms. Therefore, the ability to obtain HLA genotypes directly from scRNA-seq data represents a way to maximize the utility of these methods. We believe our study demonstrates that such direct genotyping can achieve sufficient accuracy for downstream applications unique to single-cell experiments, such as precise evaluation of ASE.

## ACKNOWLEDGEMENTS

P.K. is funded in part by the Bill and Melinda Gates Foundation (OPP1113682); the National Institute of Allergy and Infectious Diseases (NIAID) grants 1U19AI109662, U19AI057229, U19AI167903, and 5R01AI125197; Department of Defense contracts W81XWH-18-1-0253 and W81XWH-19-1-0235; and the Ralph & Marian Falk Medical Research Trust. B.D.S. is additionally funded by the National Heart, Lung, and Blood Institute (NHLBI) grant R38HL143615. J.D.G and J.R.H are funded by the National Cancer Institute grant R01CA264090.

## AUTHOR CONTRIBUTIONS

B.D.S and P.K. conceived of the study, interpreted results, and wrote the manuscript. B.D.S. and H.Z. processed raw sequencing data and B.D.S conducted subsequent data analysis. L.W.D and C.S.H. collected and analyzed previously described scRNA-seq data and performed additional sample preparation for molecular HLA genotyping of the NIH-HBM cohort. J.D.G and J.R.H. collected and analyzed previously described scRNA-seq data and provided HLA genotyping of the ISB-Swedish INCOV cohort.

---

**Supplemental Figure S1:**
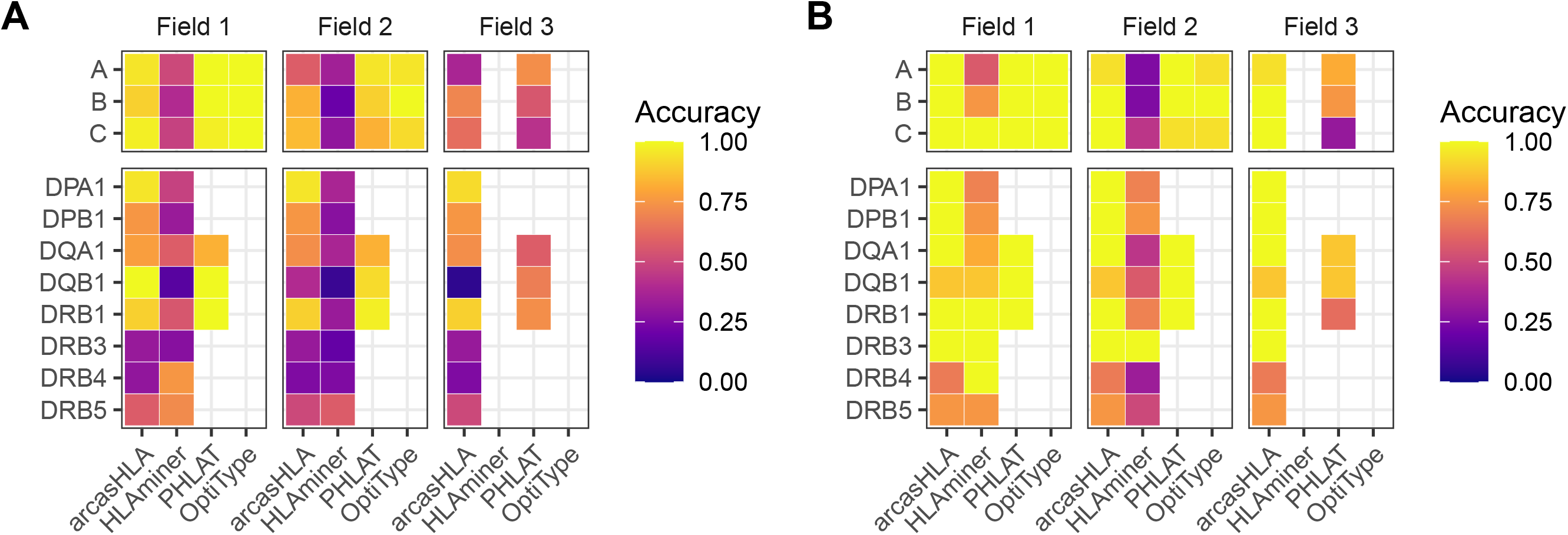
Accuracy of HLA genotype predictions from alternate sequencing platforms. Mean accuracy of predicted genotypes compared to molecular genotyping from **A)** 3’-based scRNA-seq and **B)** paired-end bulk RNA-seq data

**Supplemental Figure S2:**
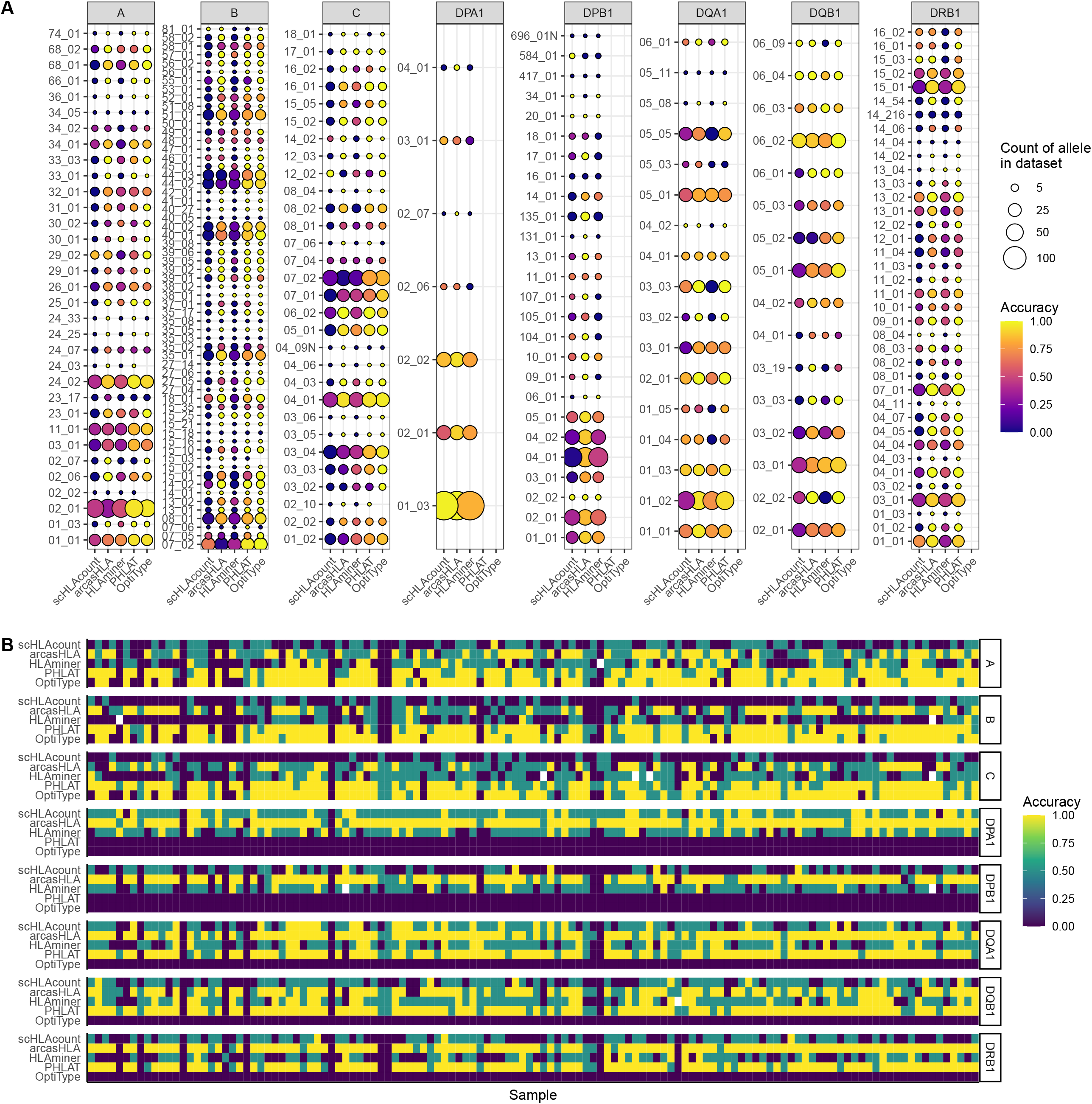
Accuracy of HLA genotype predictions by allele and sample. **A-B)** 2-field accuracy of HLA genotype predictions. **A)** Mean accuracy based on identity of true allele from molecular genotyping. Size representative of count of allele throughout entire dataset. **B)** Accuracy of genotype prediction for each.

**Supplemental Figure S3:**
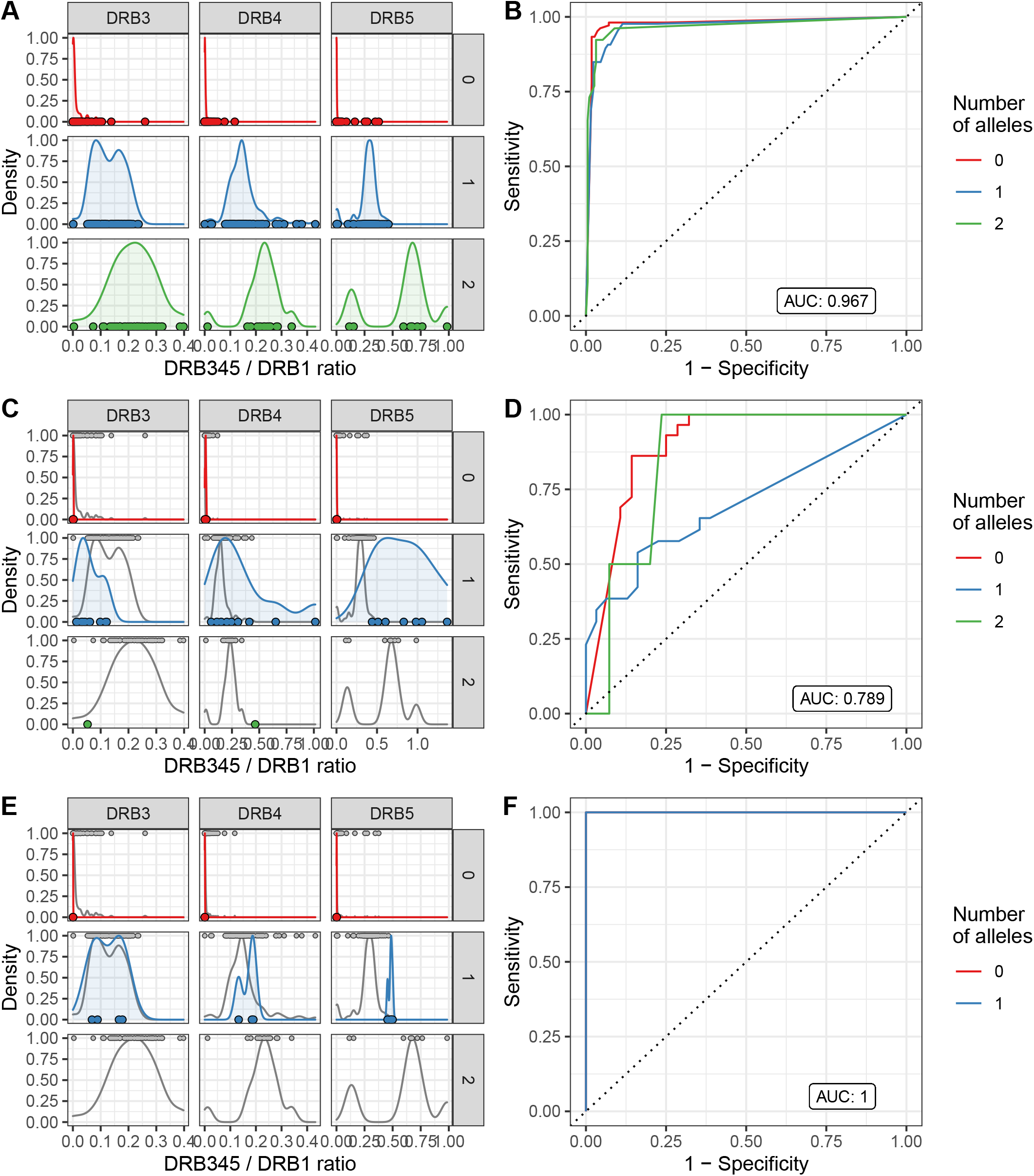
HLA-DRB345 kNN classifier performance. **A,C,E)** Input DRB345 / DRB1 read ratios and **B,D,F)** kNN model performance ROCs from **A-B)** 5’-based scRNA-seq, **C-D)** 3’-based scRNA-seq, and **E-F)** paired-end bulk RNA-seq. Distribution of HLA-DRB345 : HLA-DRB1 read ratios by locus and ground truth allele copy number used by kNN classifier. Colored distributions represent sequencing platform as indicated above. Where present, grey represents 5’-based scRNA distribution as reference. Points represent ratios from individual samples overlaid along X-axis.

**Supplemental Figure S4:**
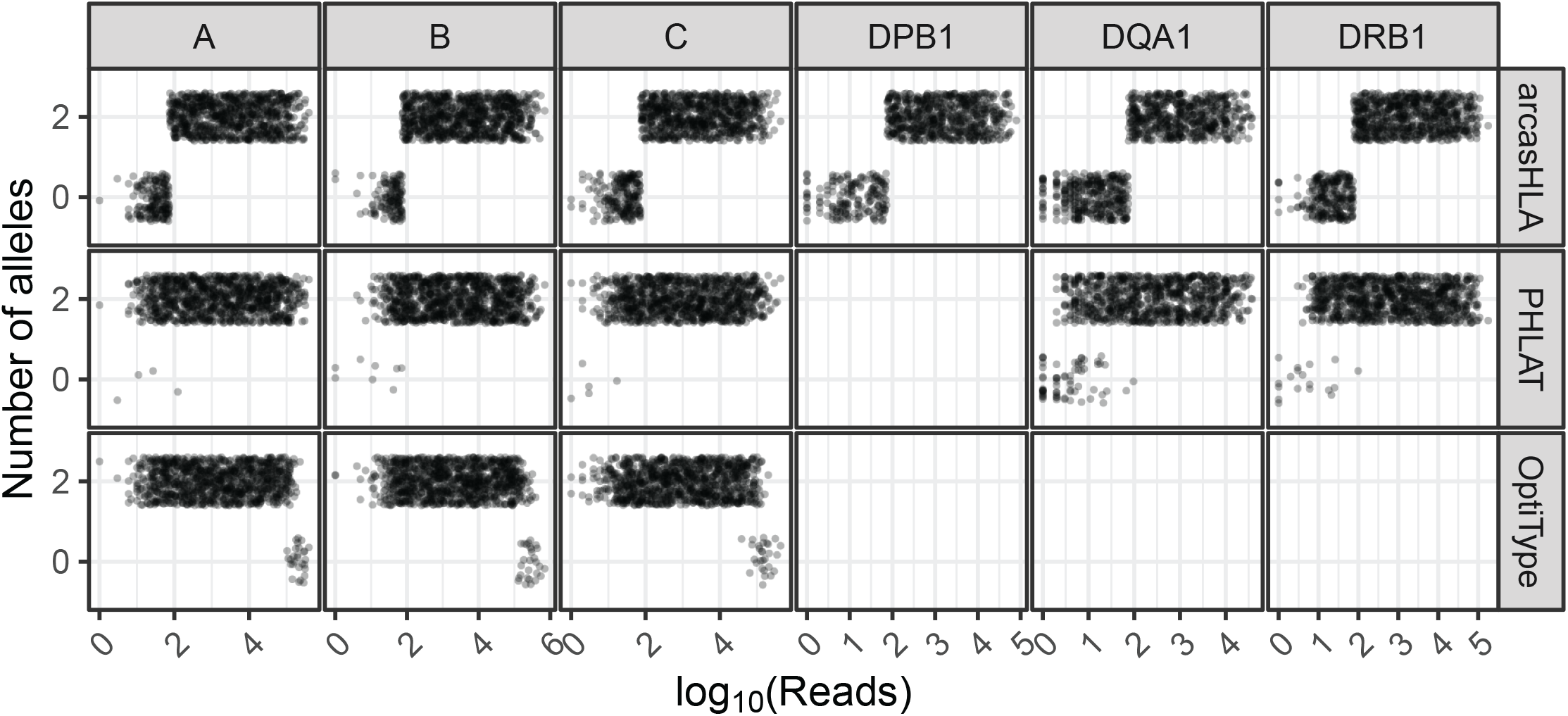
Effect of sequence sampling depth on HLA predictions. Number of alleles identified based on number of reads aligned to the indicated locus. Raw sequencing data sampled to 100%, 10%, 1%, and 0.1% of original sample read total.

**Supplemental Figure S5:**
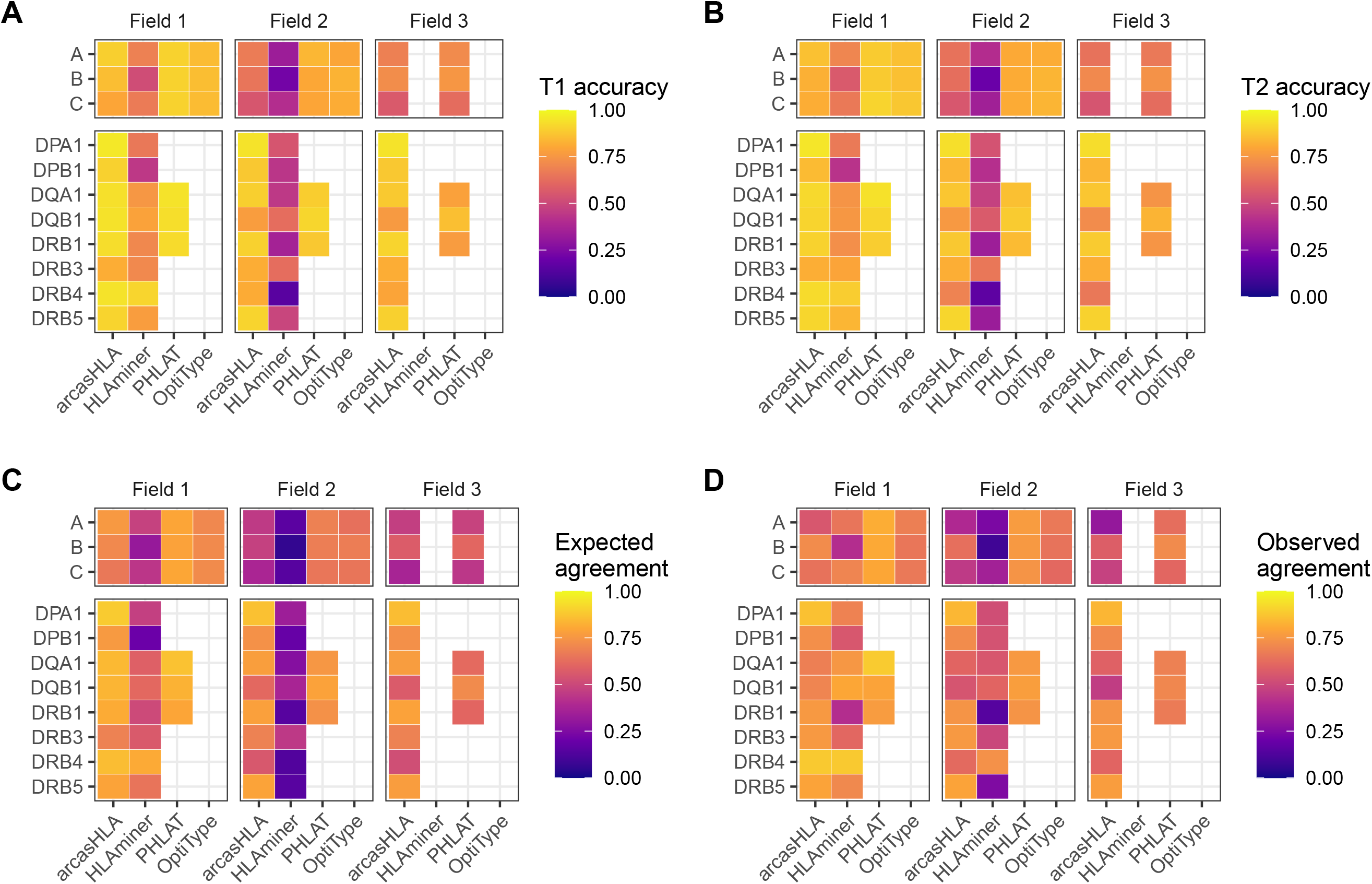
Agreement between genotype predictions at different time points. Mean accuracy of predicted genotypes compared to molecular genotyping at **A)** time point 1 and **B)** time point 2. **C)** Expected mean agreement between genotype predictions at type point 1 and type point 2 as represented by the joint probability of an accurate prediction at each time point. **D)** The observed mean agreement between predicted genotypes at time point 1 and time point 2.

**Supplemental Figure S6:**
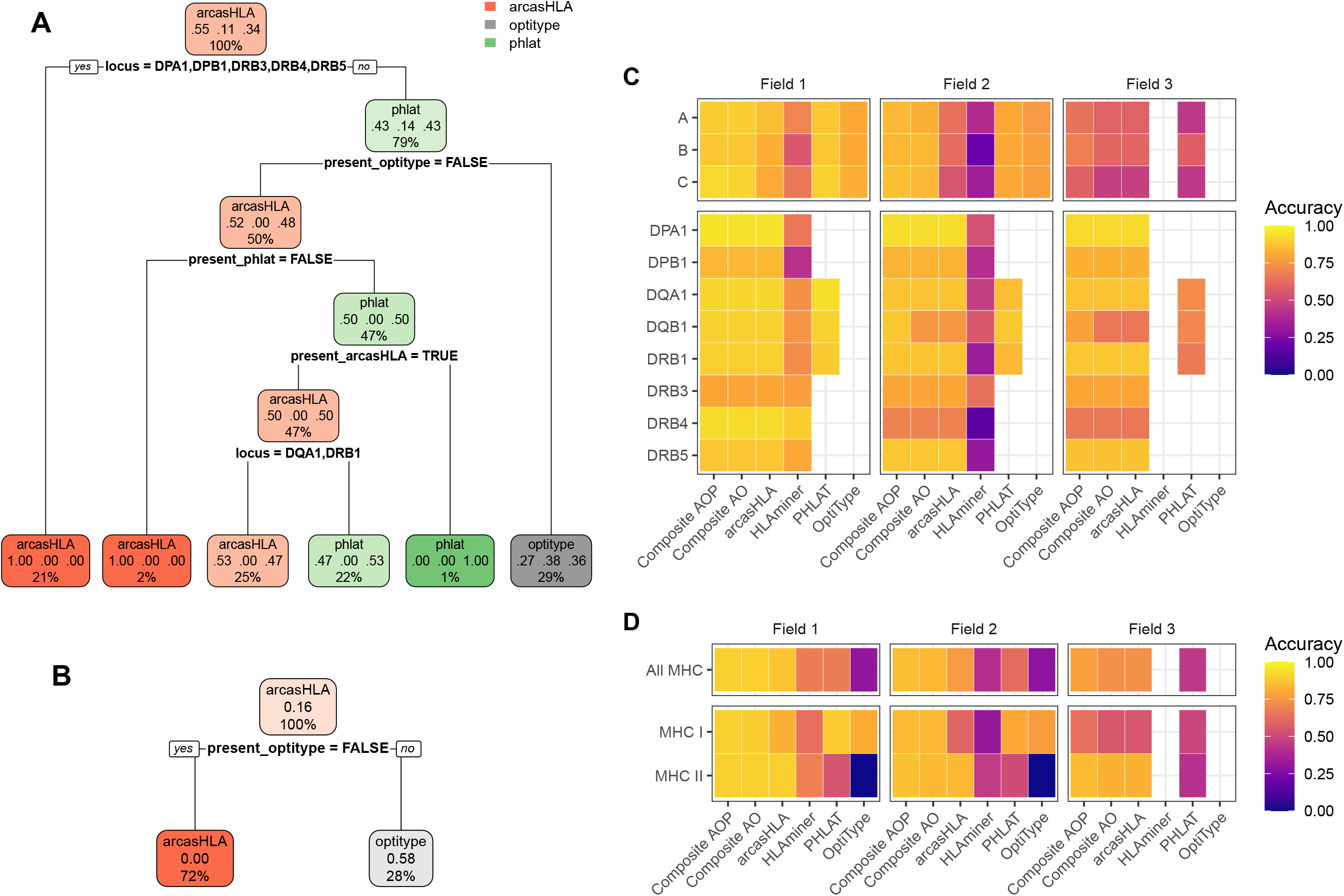
Composite accuracy. **A-B)** Decision trees for identifying the highest accuracy genotyper based the set of available genotyper predictions, locus, and field. Trained on predictions from **A)** arcasHLA, OptiType, and PHLAT or **B)** only arcasHLA and OptiType. Tuned on 70% test set with 10-fold cross validation. AUC based on performance on 30% hold-out test set. **B-C)** Mean accuracy of individual and composite genotype predictions across **B)** individual loci and **D)** loci classes.

**Supplemental Figure S7:**
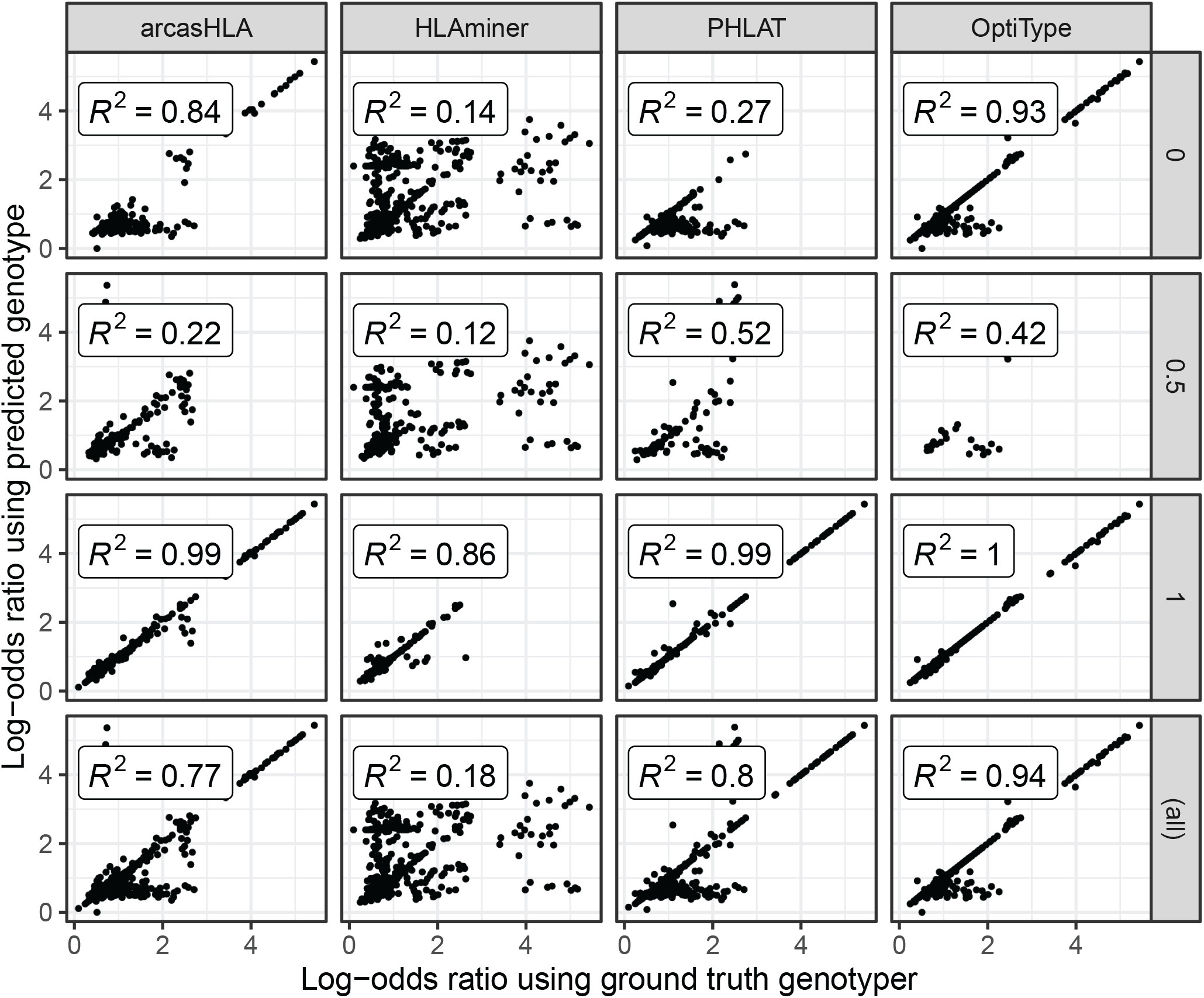
Effect of prediction accuracy on analysis of allele-specific HLA-A expression. Correlation of ground truth- and genotyper-derived HLA-A allele log-odds ratio from all samples and cell types. Rows reflect correlations stratified by underlying genotype prediction accuracy, as well as final row containing all predictions.

**Supplemental Table 1.**
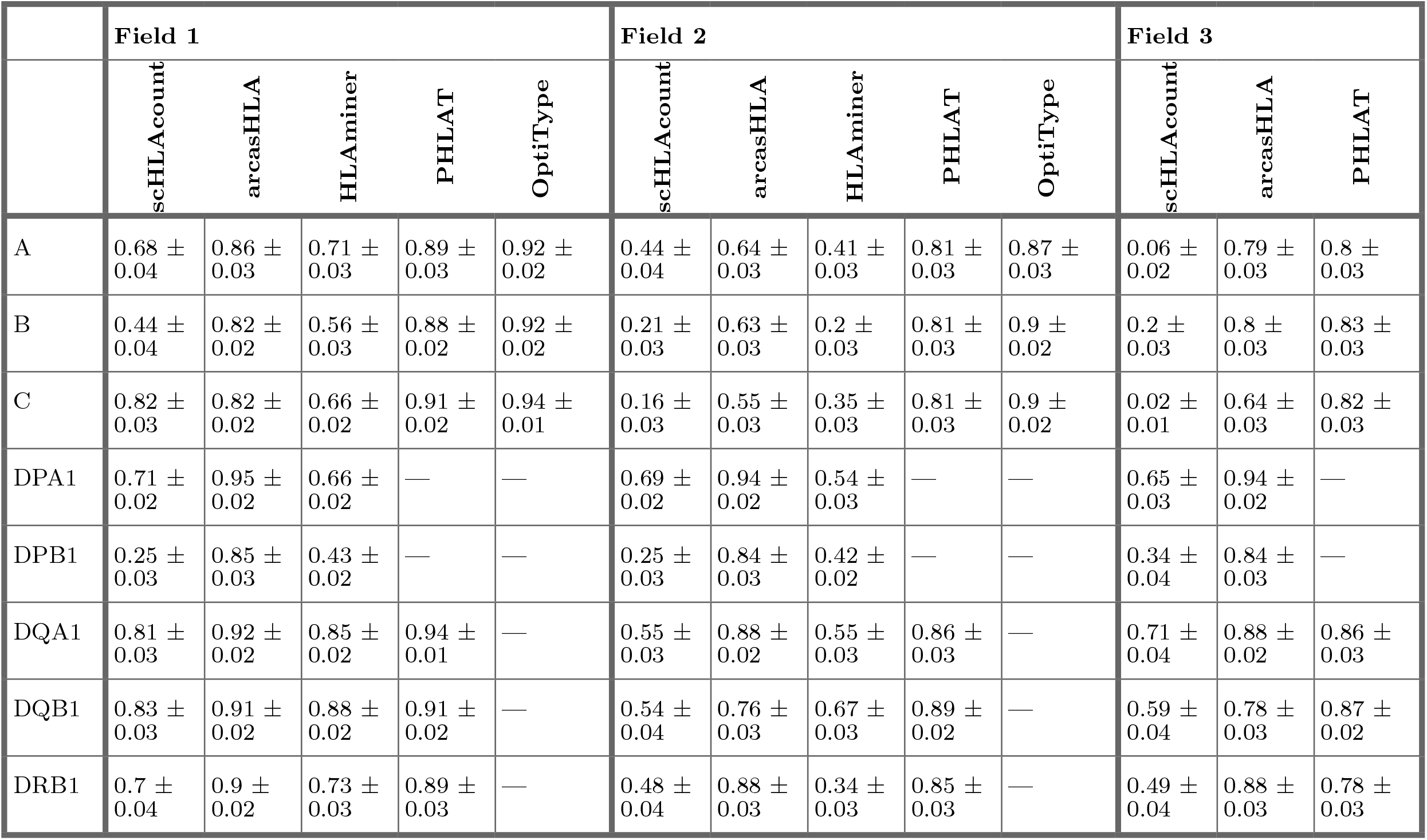
5’-based scRNA-seq success. Success of predicted genotypes from 5’-based scRNA-seq sequences. Values represent mean success +/- SEM

**Supplemental Table 2.**
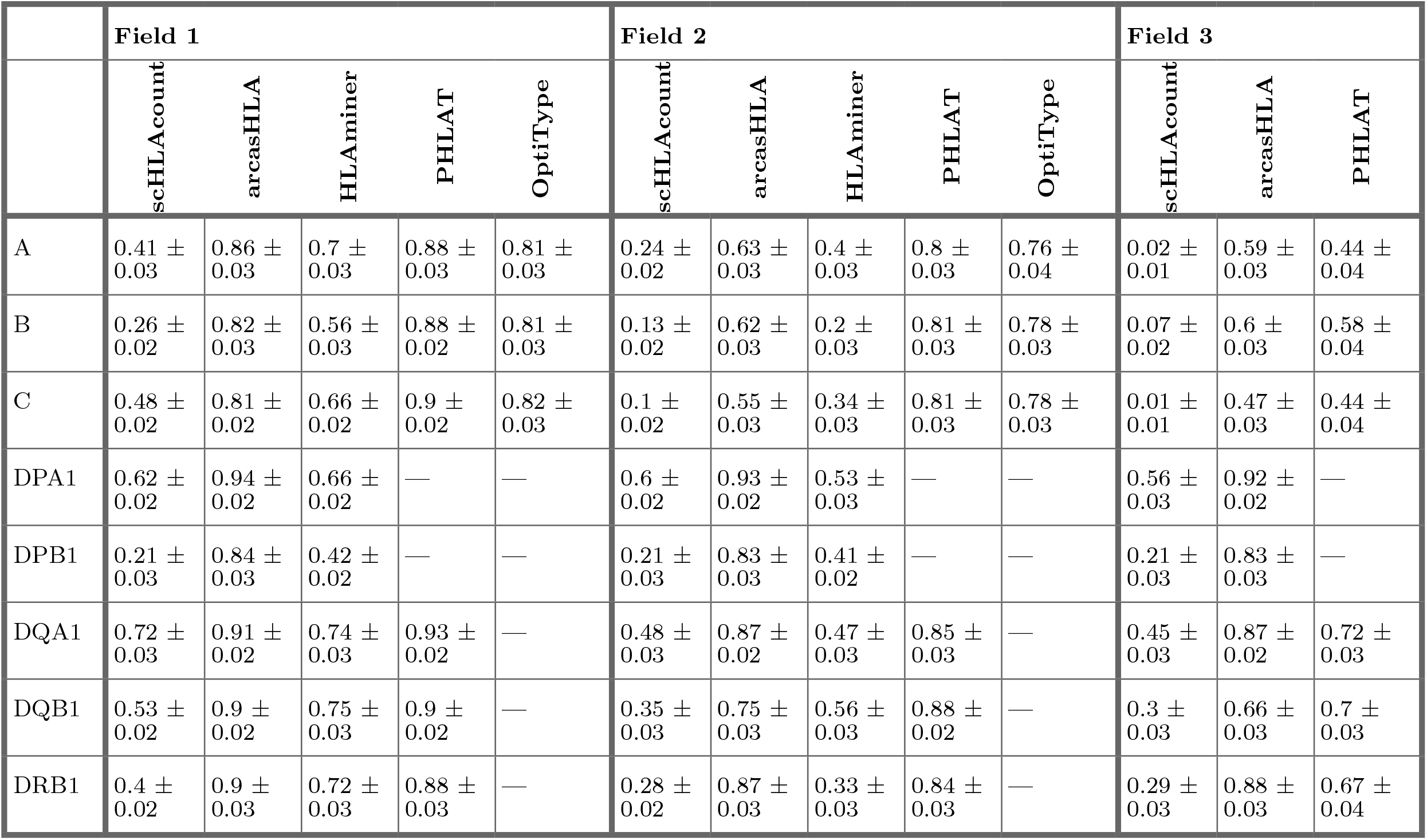
5’-based scRNA-seq accuracy. Accuracy of predicted genotypes from 5’-based scRNA-seq sequences. Values represent mean accuracy +/- SEM

**Supplemental Table 3.**
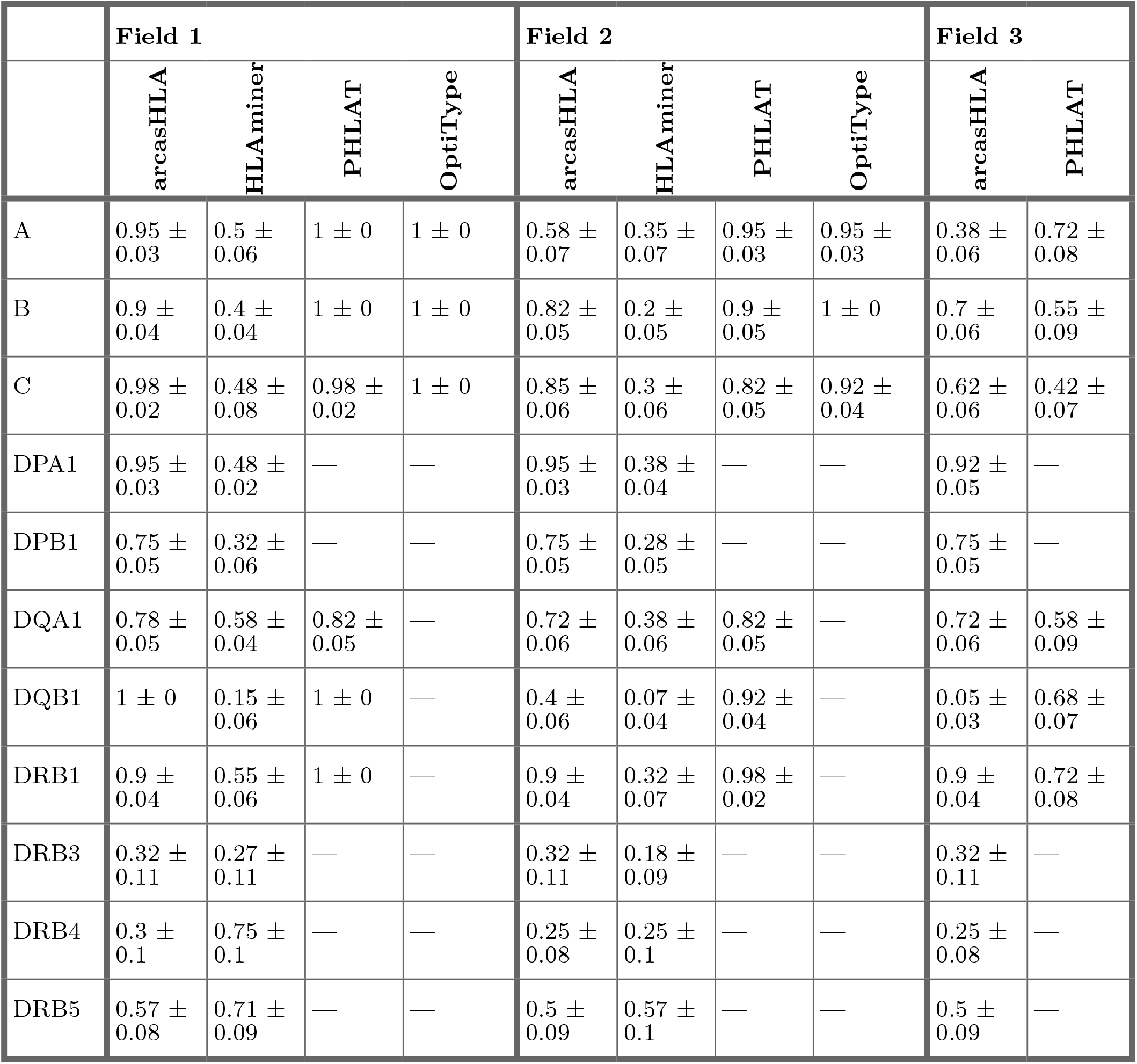
3’-based scRNA-seq accuracy. Accuracy of predicted genotypes from 3’-based scRNA-seq sequences. Values represent mean accuracy +/- SEM

|  | Field 1 |  |  |  | Field 2 |  |  |  | Field 3 |  |
| --- | --- | --- | --- | --- | --- | --- | --- | --- | --- | --- |
|  | arcasHLA | HLAminer | PHLAT | OptiType | arcasHLA | HLAminer | PHLAT | OptiType | arcasHLA | PHLAT |
| A | 0.95 ± 0.03 | 0.5 ± 0.06 | 1 ± 0 | 1 ± 0 | 0.58 ± 0.07 | 0.35 ± 0.07 | 0.95 ± 0.03 | 0.95 ± 0.03 | 0.38 ± 0.06 | 0.72 ± 0.08 |
| B | 0.9 ± 0.04 | 0.4 ± 0.04 | 1 ± 0 | 1 ± 0 | 0.82 ± 0.05 | 0.2 ± 0.05 | 0.9 ± 0.05 | 1 ± 0 | 0.7 ± 0.06 | 0.55 ± 0.09 |
| C | 0.98 ± 0.02 | 0.48 ± 0.08 | 0.98 ± 0.02 | 1 ± 0 | 0.85 ± 0.06 | 0.3 ± 0.06 | 0.82 ± 0.05 | 0.92 ± 0.04 | 0.62 ± 0.06 | 0.42 ± 0.07 |
| DPA1 | 0.95 ± 0.03 | 0.48 ± 0.02 | — | — | 0.95 ± 0.03 | 0.38 ± 0.04 | — | — | 0.92 ± 0.05 | — |
| DPB1 | 0.75 ± 0.05 | 0.32 ± 0.06 | — | — | 0.75 ± 0.05 | 0.28 ± 0.05 | — | — | 0.75 ± 0.05 | — |
| DQA1 | 0.78 ± 0.05 | 0.58 ± 0.04 | 0.82 ± 0.05 | — | 0.72 ± 0.06 | 0.38 ± 0.06 | 0.82 ± 0.05 | — | 0.72 ± 0.06 | 0.58 ± 0.09 |
| DQB1 | 1 ± 0 | 0.15 ± 0.06 | 1 ± 0 | — | 0.4 ± 0.06 | 0.07 ± 0.04 | 0.92 ± 0.04 | — | 0.05 ± 0.03 | 0.68 ± 0.07 |
| DRB1 | 0.9 ± 0.04 | 0.55 ± 0.06 | 1 ± 0 | — | 0.9 ± 0.04 | 0.32 ± 0.07 | 0.98 ± 0.02 | — | 0.9 ± 0.04 | 0.72 ± 0.08 |
| DRB3 | 0.32 ± 0.11 | 0.27 ± 0.11 | — | — | 0.32 ± 0.11 | 0.18 ± 0.09 | — | — | 0.32 ± 0.11 | — |
| DRB4 | 0.3 ± 0.1 | 0.75 ± 0.1 | — | — | 0.25 ± 0.08 | 0.25 ± 0.1 | — | — | 0.25 ± 0.08 | — |
| DRB5 | 0.57 ± 0.08 | 0.71 ± 0.09 | — | — | 0.5 ± 0.09 | 0.57 ± 0.1 | — | — | 0.5 ± 0.09 | — |

**Supplemental Table 4.** Paired end bulkRNA-seq accuracy. Accuracy of predicted genotypes from bulk RNA-seq sequences. Values represent mean accuracy +/- SEM

|  | Field 1 |  |  |  | Field 2 |  |  |  | Field 3 |  |
| --- | --- | --- | --- | --- | --- | --- | --- | --- | --- | --- |
|  | arcasHLA | HLAminer | PHLAT | OptiType | arcasHLA | HLAminer | PHLAT | OptiType | arcasHLA | PHLAT |
| A | 1 ± 0 | 0.56 ± 0.06 | 1 ± 0 | 1 ± 0 | 0.94 ± 0.06 | 0.25 ± 0.09 | 1 ± 0 | 0.94 ± 0.06 | 0.94 ± 0.06 | 0.81 ± 0.13 |
| B | 1 ± 0 | 0.75 ± 0.09 | 1 ± 0 | 1 ± 0 | 1 ± 0 | 0.25 ± 0.13 | 1 ± 0 | 1 ± 0 | 1 ± 0 | 0.75 ± 0.16 |
| C | 1 ± 0 | 1 ± 0 | 1 ± 0 | 1 ± 0 | 1 ± 0 | 0.44 ± 0.11 | 0.94 ± 0.06 | 0.94 ± 0.06 | 1 ± 0 | 0.31 ± 0.16 |
| DPA1 | 1 ± 0 | 0.69 ± 0.09 | — | — | 1 ± 0 | 0.69 ± 0.09 | — | — | 1 ± 0 | — |
| DPB1 | 1 ± 0 | 0.75 ± 0.09 | — | — | 1 ± 0 | 0.75 ± 0.09 | — | — | 1 ± 0 | — |
| DQA1 | 1 ± 0 | 0.81 ± 0.09 | 1 ± 0 | — | 1 ± 0 | 0.44 ± 0.15 | 1 ± 0 | — | 1 ± 0 | 0.88 ± 0.08 |
| DQB1 | 0.88 ± 0.08 | 0.88 ± 0.08 | 1 ± 0 | — | 0.88 ± 0.08 | 0.56 ± 0.11 | 1 ± 0 | — | 0.88 ± 0.08 | 0.88 ± 0.08 |
| DRB1 | 1 ± 0 | 1 ± 0 | 1 ± 0 | — | 1 ± 0 | 0.69 ± 0.09 | 1 ± 0 | — | 1 ± 0 | 0.62 ± 0.16 |
| DRB3 | 1 ± 0 | 1 ± 0 | — | — | 1 ± 0 | 1 ± 0 | — | — | 1 ± 0 | — |
| DRB4 | 0.67 ± 0.1 | 1 ± 0 | — | — | 0.67 ± 0.1 | 0.33 ± 0.2 | — | — | 0.67 ± 0.1 | — |
| DRB5 | 0.75 ± 0.18 | 0.75 ± 0.18 | — | — | 0.75 ± 0.18 | 0.5 ± 0.2 | — | — | 0.75 ± 0.18 | — |

**Supplemental Table 5.**
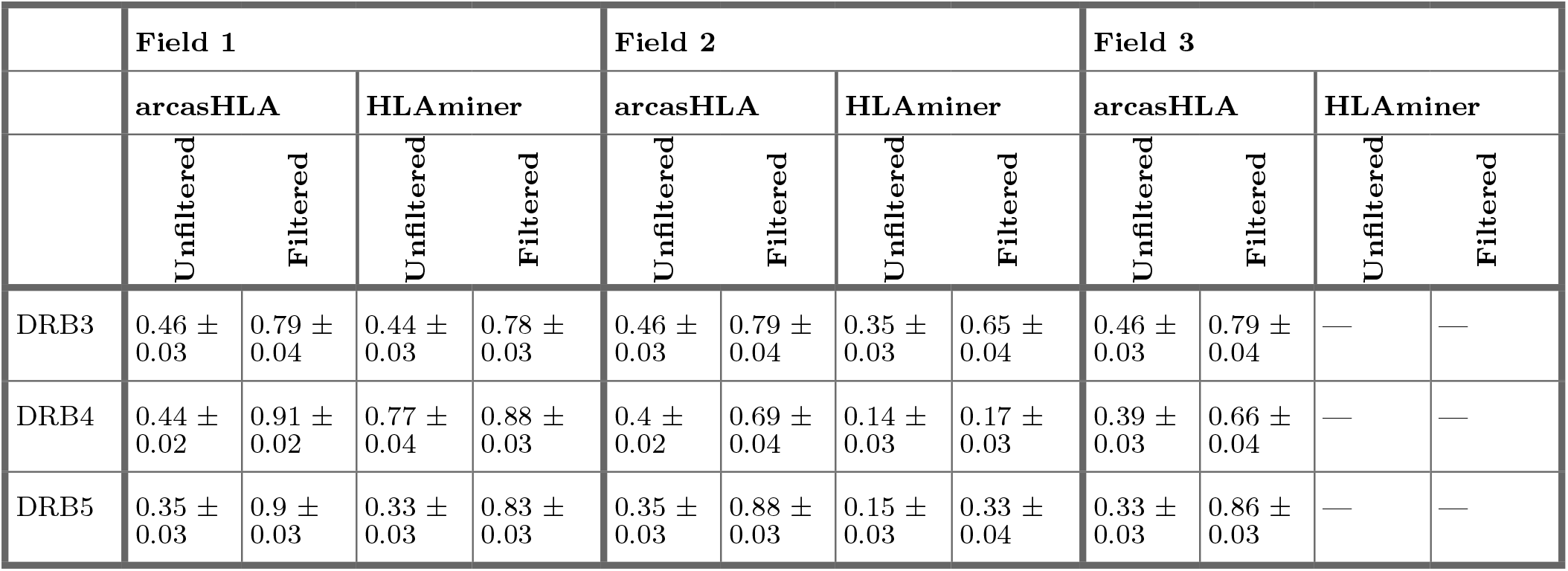
*HLA-DRB345* accuracy. Accuracy of *HLA-DRB345* genotypes filtered or unfiltered by kNN copy number classifier. Values represent mean accuracy +/- SEM

**Supplemental Table 6.**
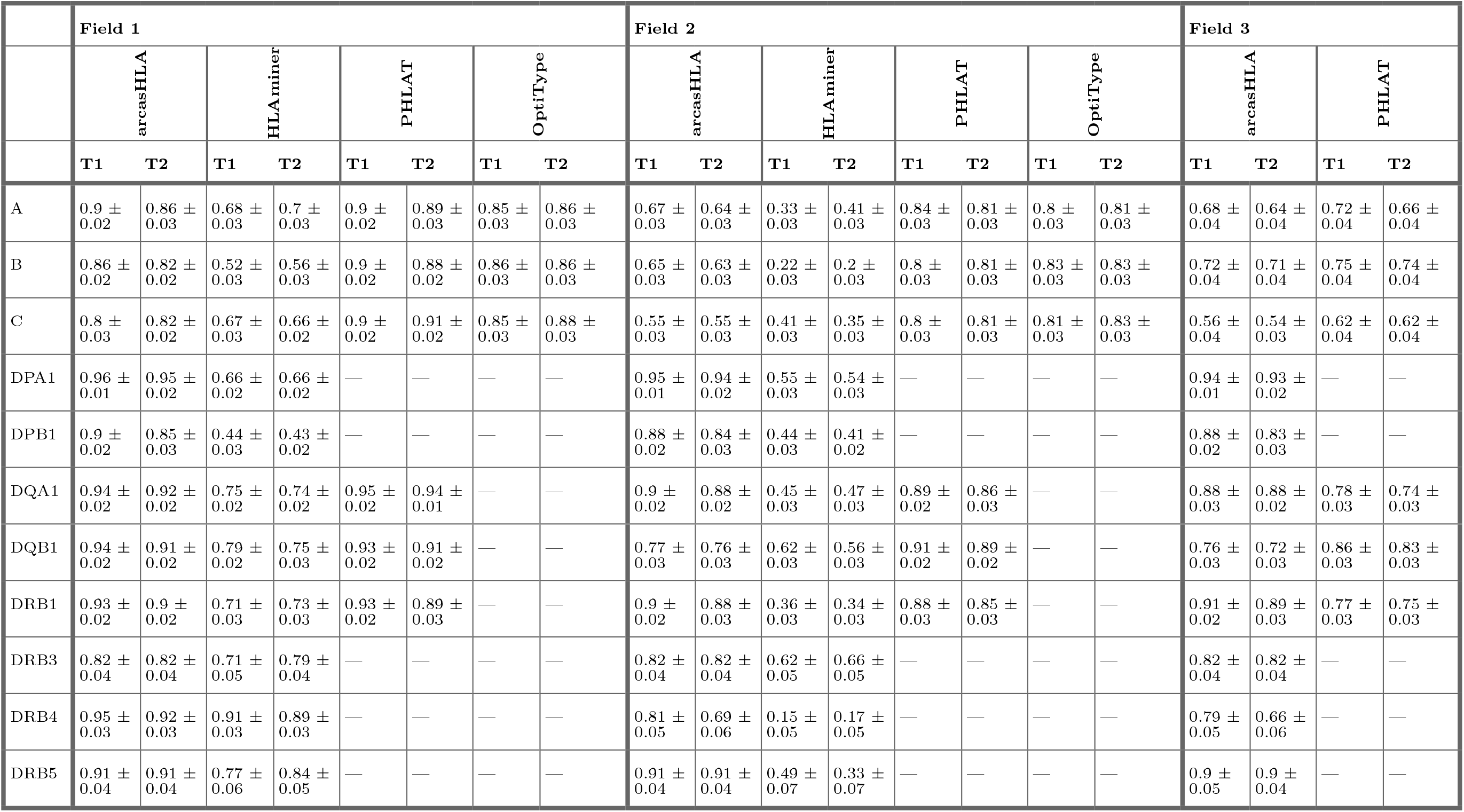
Reproducibility of genotype prediction accuracy. Accuracy of predicted genotypes from repeated sampling of individuals at time point 1 (T1) and time point 2 (T2). Values represent mean accuracy +/- SEM

|  | Field 1 |  |  |  |  |  |  |  | Field 2 |  |  |  |  |  |  |  | Field 3 |  |  |  |
| --- | --- | --- | --- | --- | --- | --- | --- | --- | --- | --- | --- | --- | --- | --- | --- | --- | --- | --- | --- | --- |
|  | arcasHLA |  | HLAminer |  | PHLAT |  | OptiType |  | arcasHLA |  | HLAminer |  | PHLAT |  | OptiType |  | arcasHLA |  | PHLAT |  |
|  | T1 | T2 | T1 | T2 | T1 | T2 | T1 | T2 | T1 | T2 | T1 | T2 | T1 | T2 | T1 | T2 | T1 | T2 | T1 | T2 |
| A | 0.9 ± 0.02 | 0.86 ± 0.03 | 0.68 ± 0.03 | 0.7 ± 0.03 | 0.9 ± 0.02 | 0.89 ± 0.03 | 0.85 ± 0.03 | 0.86 ± 0.03 | 0.67 ± 0.03 | 0.64 ± 0.03 | 0.33 ± 0.03 | 0.41 ± 0.03 | 0.84 ± 0.03 | 0.81 ± 0.03 | 0.8 ± 0.03 | 0.81 ± 0.03 | 0.68 ± 0.04 | 0.64 ± 0.04 | 0.72 ± 0.04 | 0.66 ± 0.04 |
| B | 0.86 ± 0.02 | 0.82 ± 0.02 | 0.52 ± 0.03 | 0.56 ± 0.03 | 0.9 ± 0.02 | 0.88 ± 0.02 | 0.86 ± 0.03 | 0.86 ± 0.03 | 0.65 ± 0.03 | 0.63 ± 0.03 | 0.22 ± 0.03 | 0.2 ± 0.03 | 0.8 ± 0.03 | 0.81 ± 0.03 | 0.83 ± 0.03 | 0.83 ± 0.03 | 0.72 ± 0.04 | 0.71 ± 0.04 | 0.75 ± 0.04 | 0.74 ± 0.04 |
| C | 0.8 ± 0.03 | 0.82 ± 0.02 | 0.67 ± 0.03 | 0.66 ± 0.02 | 0.9 ± 0.02 | 0.91 ± 0.02 | 0.85 ± 0.03 | 0.88 ± 0.03 | 0.55 ± 0.03 | 0.55 ± 0.03 | 0.41 ± 0.03 | 0.35 ± 0.03 | 0.8 ± 0.03 | 0.81 ± 0.03 | 0.81 ± 0.03 | 0.83 ± 0.03 | 0.56 ± 0.04 | 0.54 ± 0.03 | 0.62 ± 0.04 | 0.62 ± 0.04 |
| DPA1 | 0.96 ± 0.01 | 0.95 ± 0.02 | 0.66 ± 0.02 | 0.66 ± 0.02 | — | — | — | — | 0.95 ± 0.01 | 0.94 ± 0.02 | 0.55 ± 0.03 | 0.54 ± 0.03 | — | — | — | — | 0.94 ± 0.01 | 0.93 ± 0.02 | — | — |
| DPB1 | 0.9 ± 0.02 | 0.85 ± 0.03 | 0.44 ± 0.03 | 0.43 ± 0.02 | — | — | — | — | 0.88 ± 0.02 | 0.84 ± 0.03 | 0.44 ± 0.03 | 0.41 ± 0.02 | — | — | — | — | 0.88 ± 0.02 | 0.83 ± 0.03 | — | — |
| DQA1 | 0.94 ± 0.02 | 0.92 ± 0.02 | 0.75 ± 0.02 | 0.74 ± 0.02 | 0.95 ± 0.02 | 0.94 ± 0.01 | — | — | 0.9 ± 0.02 | 0.88 ± 0.02 | 0.45 ± 0.03 | 0.47 ± 0.03 | 0.89 ± 0.02 | 0.86 ± 0.03 | — | — | 0.88 ± 0.03 | 0.88 ± 0.02 | 0.78 ± 0.03 | 0.74 ± 0.03 |
| DQB1 | 0.94 ± 0.02 | 0.91 ± 0.02 | 0.79 ± 0.02 | 0.75 ± 0.03 | 0.93 ± 0.02 | 0.91 ± 0.02 | — | — | 0.77 ± 0.03 | 0.76 ± 0.03 | 0.62 ± 0.03 | 0.56 ± 0.03 | 0.91 ± 0.02 | 0.89 ± 0.02 | — | — | 0.76 ± 0.03 | 0.72 ± 0.03 | 0.86 ± 0.03 | 0.83 ± 0.03 |
| DRB1 | 0.93 ± 0.02 | 0.9 ± 0.02 | 0.71 ± 0.03 | 0.73 ± 0.03 | 0.93 ± 0.02 | 0.89 ± 0.03 | — | — | 0.9 ± 0.02 | 0.88 ± 0.03 | 0.36 ± 0.03 | 0.34 ± 0.03 | 0.88 ± 0.03 | 0.85 ± 0.03 | — | — | 0.91 ± 0.02 | 0.89 ± 0.03 | 0.77 ± 0.03 | 0.75 ± 0.03 |
| DRB3 | 0.82 ± 0.04 | 0.82 ± 0.04 | 0.71 ± 0.05 | 0.79 ± 0.04 | — | — | — | — | 0.82 ± 0.04 | 0.82 ± 0.04 | 0.62 ± 0.05 | 0.66 ± 0.05 | — | — | — | — | 0.82 ± 0.04 | 0.82 ± 0.04 | — | — |
| DRB4 | 0.95 ± 0.03 | 0.92 ± 0.03 | 0.91 ± 0.03 | 0.89 ± 0.03 | — | — | — | — | 0.81 ± 0.05 | 0.69 ± 0.06 | 0.15 ± 0.05 | 0.17 ± 0.05 | — | — | — | — | 0.79 ± 0.05 | 0.66 ± 0.06 | — | — |
| DRB5 | 0.91 ± 0.04 | 0.91 ± 0.04 | 0.77 ± 0.06 | 0.84 ± 0.05 | — | — | — | — | 0.91 ± 0.04 | 0.91 ± 0.04 | 0.49 ± 0.07 | 0.33 ± 0.07 | — | — | — | — | 0.9 ± 0.05 | 0.9 ± 0.04 | — | — |

**Supplemental Table 7.** Composite genotype accuracy. Success of predicted genotypes from using decision tree-based composites. AOP = arcasHLA, OptiType, and PHLAT, AO = arcasHLA and OptiType. Values represent mean accuracy +/- SEM

|  | Field 1 |  |  |  |  |  | Field 2 |  |  |  |  |  | Field 3 |  |  |  |
| --- | --- | --- | --- | --- | --- | --- | --- | --- | --- | --- | --- | --- | --- | --- | --- | --- |
|  | Composite AOP | Composite AO | arcasHLA | HLAminer | PHLAT | OptiType | Composite AOP | Composite AO | arcasHLA | HLAminer | PHLAT | OptiType | Composite AOP | Composite AO | arcasHLA | PHLAT |
| MHC All | 0.9 ± 0.02 | 0.89 ± 0.02 | 0.87 ± 0.02 | 0.67 ± 0.01 | 0.67 ± 0.01 | 0.3 ± 0.01 | 0.86 ± 0.02 | 0.84 ± 0.02 | 0.76 ± 0.02 | 0.41 ± 0.01 | 0.62 ± 0.02 | 0.29 ± 0.01 | 0.78 ± 0.02 | 0.74 ± 0.02 | 0.74 ± 0.02 | 0.45 ± 0.02 |
| MHC I | 0.9 ± 0.02 | 0.89 ± 0.02 | 0.83 ± 0.02 | 0.63 ± 0.02 | 0.89 ± 0.02 | 0.81 ± 0.03 | 0.85 ± 0.02 | 0.84 ± 0.02 | 0.6 ± 0.02 | 0.31 ± 0.02 | 0.8 ± 0.02 | 0.78 ± 0.03 | 0.64 ± 0.02 | 0.56 ± 0.02 | 0.56 ± 0.02 | 0.49 ± 0.03 |
| MHC II | 0.9 ± 0.02 | 0.9 ± 0.02 | 0.9 ± 0.02 | 0.68 ± 0.02 | 0.54 ± 0.01 | 0 ± 0 | 0.87 ± 0.02 | 0.84 ± 0.02 | 0.84 ± 0.02 | 0.46 ± 0.02 | 0.52 ± 0.01 | 0 ± 0 | 0.85 ± 0.02 | 0.83 ± 0.02 | 0.83 ± 0.02 | 0.42 ± 0.01 |
| A | 0.89 ± 0.03 | 0.89 ± 0.03 | 0.86 ± 0.03 | 0.7 ± 0.03 | 0.88 ± 0.03 | 0.81 ± 0.03 | 0.84 ± 0.03 | 0.82 ± 0.03 | 0.63 ± 0.03 | 0.4 ± 0.03 | 0.8 ± 0.03 | 0.76 ± 0.04 | 0.64 ± 0.03 | 0.59 ± 0.03 | 0.59 ± 0.03 | 0.44 ± 0.04 |
| B | 0.88 ± 0.02 | 0.88 ± 0.02 | 0.82 ± 0.03 | 0.56 ± 0.03 | 0.88 ± 0.02 | 0.81 ± 0.03 | 0.84 ± 0.03 | 0.85 ± 0.03 | 0.62 ± 0.03 | 0.2 ± 0.03 | 0.81 ± 0.03 | 0.78 ± 0.03 | 0.68 ± 0.03 | 0.6 ± 0.03 | 0.6 ± 0.03 | 0.58 ± 0.04 |
| C | 0.92 ± 0.02 | 0.91 ± 0.02 | 0.81 ± 0.02 | 0.66 ± 0.02 | 0.9 ± 0.02 | 0.82 ± 0.03 | 0.86 ± 0.02 | 0.84 ± 0.02 | 0.55 ± 0.03 | 0.34 ± 0.03 | 0.81 ± 0.03 | 0.78 ± 0.03 | 0.59 ± 0.03 | 0.47 ± 0.03 | 0.47 ± 0.03 | 0.44 ± 0.04 |
| DPA1 | 0.94 ± 0.02 | 0.94 ± 0.02 | 0.94 ± 0.02 | 0.66 ± 0.02 | — | — | 0.93 ± 0.02 | 0.93 ± 0.02 | 0.93 ± 0.02 | 0.53 ± 0.03 | — | — | 0.92 ± 0.02 | 0.92 ± 0.02 | 0.92 ± 0.02 | — |
| DPB1 | 0.84 ± 0.03 | 0.84 ± 0.03 | 0.84 ± 0.03 | 0.42 ± 0.02 | — | — | 0.83 ± 0.03 | 0.83 ± 0.03 | 0.83 ± 0.03 | 0.41 ± 0.02 | — | — | 0.83 ± 0.03 | 0.83 ± 0.03 | 0.83 ± 0.03 | — |
| DQA1 | 0.91 ± 0.02 | 0.91 ± 0.02 | 0.91 ± 0.02 | 0.74 ± 0.03 | 0.93 ± 0.02 | — | 0.87 ± 0.02 | 0.87 ± 0.02 | 0.87 ± 0.02 | 0.47 ± 0.03 | 0.85 ± 0.03 | — | 0.87 ± 0.02 | 0.87 ± 0.02 | 0.87 ± 0.02 | 0.72 ± 0.03 |
| DQB1 | 0.9 ± 0.02 | 0.9 ± 0.02 | 0.9 ± 0.02 | 0.75 ± 0.03 | 0.9 ± 0.02 | — | 0.88 ± 0.02 | 0.75 ± 0.03 | 0.75 ± 0.03 | 0.56 ± 0.03 | 0.88 ± 0.02 | — | 0.79 ± 0.03 | 0.66 ± 0.03 | 0.66 ± 0.03 | 0.7 ± 0.03 |
| DRB1 | 0.9 ± 0.03 | 0.9 ± 0.03 | 0.9 ± 0.03 | 0.72 ± 0.03 | 0.88 ± 0.03 | — | 0.87 ± 0.03 | 0.87 ± 0.03 | 0.87 ± 0.03 | 0.33 ± 0.03 | 0.84 ± 0.03 | — | 0.88 ± 0.03 | 0.88 ± 0.03 | 0.88 ± 0.03 | 0.67 ± 0.04 |
| DRB3 | 0.8 ± 0.04 | 0.8 ± 0.04 | 0.8 ± 0.04 | 0.77 ± 0.03 | — | — | 0.8 ± 0.04 | 0.8 ± 0.04 | 0.8 ± 0.04 | 0.64 ± 0.04 | — | — | 0.8 ± 0.04 | 0.8 ± 0.04 | 0.8 ± 0.04 | — |
| DRB4 | 0.92 ± 0.02 | 0.92 ± 0.02 | 0.92 ± 0.02 | 0.89 ± 0.02 | — | — | 0.69 ± 0.04 | 0.69 ± 0.04 | 0.69 ± 0.04 | 0.17 ± 0.03 | — | — | 0.66 ± 0.04 | 0.66 ± 0.04 | 0.66 ± 0.04 | — |
| DRB5 | 0.88 ± 0.03 | 0.88 ± 0.03 | 0.88 ± 0.03 | 0.8 ± 0.03 | — | — | 0.88 ± 0.03 | 0.88 ± 0.03 | 0.88 ± 0.03 | 0.31 ± 0.04 | — | — | 0.86 ± 0.03 | 0.86 ± 0.03 | 0.86 ± 0.03 | — |

